# PSMA-bearing extracellular vesicles secreted from prostate cancer convert the microenvironment to a tumor-supporting, pro-angiogenic state

**DOI:** 10.1101/2022.02.25.482024

**Authors:** Camila Maria Longo Machado, Magdalena Skubal, Katja Haedicke, Fabio Pittella Silva, Evan Paul Stater, Thais Larissa Araujo de Oliveira Silva, Erico Tosoni Costa, Cibele Masotti, Andreia Hanada Otake, Luciana Nogueira Sousa Andrade, Mara de Souza Junqueira, Hsiao-Ting Hsu, Sudeep Das, Benedict Mc Larney, Edwin Charles Pratt, Yevgeniy Romin, Ning Fan, Katia Manova-Todorova, Martin Pomper, Jan Grimm

## Abstract

Extracellular vesicles (EV) are comprised of vesicles budding from cell membranes and smaller intracellular vesicles shed by cells. EV play a role in remodeling the tumor microenvironment (TME) and support tumor progression. Prostate-specific membrane antigen (PSMA) is a transmembrane glycoprotein with a carboxypeptidase function, frequently associated with poor clinical prognosis in prostate cancer (PCa). We previously identified an oncogenic PSMA signaling function in prostate cancer. Others demonstrated that EV isolated from the plasma of patients with high-grade PCa carry PSMA, but so far no pathophysiological effect has been associated with PSMA-bearing EV. Here we demonstrate that EV from PCa cells are able to transfer PSMA and its functionality to cells in the TME. The consequence of that EV-mediated PSMA transfer is an acute to long-term increased secretion of vascular endothelial growth factor-A (VEGF-A), angiogenin, pro-angiogenic and pro-lymphangiogenic mediators and increased 4E binding protein 1 (4EBP-1) phosphorylation in tumors. We compare EV from PCa cells with or without PSMA expression to address the role of PSMA-bearing EV in promoting pro-tumoral changes in the TME using classical molecular biology and novel molecular imaging approaches.

## Introduction

Extracellular vesicles (EV) include apoptotic bodies, microvesicles, and exosomes. They are sub-categorized by biogenesis and size (Ailuno et al., 2020). Apoptotic bodies are generated by membrane disintegration after injury and cell death, producing vesicles ranging from 1 - 5 µm in diameter. Microvesicles are produced from viable cells by the plasma membrane bulging outward, ranging from 50 - 1,000 nm in diameter (Ailuno et al., 2020).Exosomes are the smallest and single membrane units shed by cells, and loaded with intracellular organelles and contents, with a diameter ranging from 20 to 160 nm (Ailuno et al., 2020). EV from prostate cancer (PCa) cells carry prostate-specific membrane antigen (PSMA), a transmembrane glycoprotein with carboxypeptidase function. PSMA is highly expressed in primary PCa, recurrent tumors, and in regional and distant metastases (Biggs et al., 2016; Hupe et al., 2018; Watanabe et al., 2021). Studies have explored EV enriched with PSMA as a diagnostic tool without further investigating their biological role in detail (Biggs et al., 2016; Joncas et al., 2019; Krishn et al., 2019; Liu et al., 2014; Minciacchi et al., 2017; Mitchell et al., 2009; Notarangelo et al., 2019; Park et al., 2016). In patients with PCa, several subtypes of EV were identified (Turay et al., 2016) with diameters ranging between 100 nm - 1 µm (Minciacchi et al., 2015). EV preserve the donor cell membrane’s identity, including the milieu of biomolecules (Di Vizio et al., 2012). Several studies have shown that patients with higher Gleason scores have a correspondingly higher number of PSMA-carrying EV present in plasma (Padda et al., 2019). Padda et al. (Padda et al., 2019) and Krishn et al. (Krishn et al., 2019) characterized EV from patient plasma as a subtype with a size range from 100 nm - 1 µm that were enriched in PSMA, CD9-tetraspanin, and β1-integrin (Bordas et al., 2020; Kowal et al., 2016). A subsequent study (Joncas et al., 2019) isolated a mixed population of EV with a mean size of 200 nm from patients with metastatic PCa. EV have emerged as a means to modify the TME to establish a pre-metastatic niche (Padda et al., 2019) and lymph nodes (LN) metastases (García-Silva et al., 2021).

Tumor angiogenesis is assisted by factors secreted by cells within the TME to recruit stroma cells and develop new blood vessels (Antonarakis and Carducci, 2012). Vascular endothelial growth factor-A (VEGF-A) and angiogenin are early pro-angiogenic factors highly expressed in the TME (Liu et al., 2015) and in sites of inflammation (Barrientos et al., 2008). Weidner et al. showed that increased microvessel density in primary cancer samples (Weidner et al., 1993) is associated with poor prognosis (Liu et al., 2015; Pore et al., 2003; Senthil et al., 2003; Taraboletti et al., 2006; Zhan et al., 2013). High VEGF-A levels in human and murine PCa samples have been associated with poor survival and tumor recurrence (Duque et al., 1999; Ferrara, 2002; Liu et al., 2015; Picus et al., 2011; Ungersma et al., 2010). VEGF-A can also act as a secondary messenger to recruit tumor-associated macrophages (TAM) to solid tumors (Lewis et al., 2000). It has been shown that Ak strain transforming (AKT) activation (Ghosh et al., 2010) up-regulates VEGF while AKT phosphorylation-inhibition downregulates VEGF-A expression (Pore et al., 2006). We have previously shown a positive correlation between PSMA expression and poor prognosis of PCa in tissues with increased detection of phosphor-4EBP-1 (Kaittanis et al., 2018). 4EBP-1 phosphorylation is a downstream marker of AKT-pathway activation in tumor tissues (Kaittanis et al., 2018) and associated with increased VEGF expression independently of hypoxic events (De Benedetti and Graff, 2004). Here we show that circulating EV derived from PCa-PSMA-expressing cells (PSMA-EV) can induce PSMA-negative cells to secrete VEGF, angiogenin and pro-lymphangiogenic factors upon accumulation of PSMA. Specifically, we observed that PSMA-EV increased VEGF-A and angiogenin secretion from both tumor and endothelial cells. Those two factors enhanced microvascular density in the TME and initiated a lymphangiogenic switch through increased PDGF-BB, PlGF, ANG-2, CD68^+^, podoplanin^+^, and PDGFRβ1^+^. Our data indicate that tumor cells directly influence their environment through secreted EV that includes biologically active components, with potentially far-reaching consequences for tumor development, detection and therapy.

## Results

### PSMA-EV transfer biologically active PSMA to recipient cells

To investigate whether PSMA-EV cargo triggers pro-angiogenic factors in PCa, we utilized the PSMA-negative human PCa cell line PC3 (PC3WT) as a recipient of EV from PSMA expressing cells. We established an EV isolation method and characterized isolated EV by determining EV marker expression and assaying the potential of isolated EV to transfer detectable PSMA to recipient cells. Nuanced differences in size distribution were observed among the EV isolated from different PSMA-PCa donor cell lines. Characterization of isolated EV, and *in vitro* functional assays of EV uptake and cargo transfer to recipient cells are shown in the supplementary data (Supplementary Fig. 1 and 2).

Our results showed that overexpression of PSMA in PC3 (PC3PSMA) did not affect the size of the shed EV (PSMA-EV) (Fig. 1A, Supplementary Fig. 1A-D), consistent with previously published results for EV derived from PC3 wild type (WT) cells (WT-EV) (Smyth et al., 2015). Isolated WT-EV and PSMA-EV were of consistent size across multiple isolation batches (Fig. 1B, Supplementary Fig. 1D). The isolation method established here (Supplementary Fig. 1A) selected for EV, comparable to traditional isolation methods such as ultracentrifugation followed by a sucrose density cushion (Bordas et al., 2020; Gupta et al., 2020; Kowal et al., 2016; Lázaro-Ibáñez et al., 2017; Padda et al., 2019; Smyth et al., 2015; Todorova et al., 2017; Zaborowski et al., 2015). Immunoblotting of total extracted EV protein showed EV markers (flotilin-1, CD9, β1-integrin, and β-actin) carried by the various EV types (Fig. 1C). Immunoblotting confirmed that PSMA-EV (but not WT-EV) expressed PSMA (Fig.1C). Furthermore, PSMA transference was detectable from PSMA-EV to recipient cells (Supplementary Fig. 1G – 2D). PSMA-EV induced PSMA expression could be observed for up to 48 h in PC3 WT cells without cellular toxicity or morphological alterations. PSMA-EV and WT-EV were taken up at similar rates by recipient cells (Fig.1D, Supplementary Fig. 1M to 1P and Supplementary Fig. 2A to 2D).

**Figure 1:**
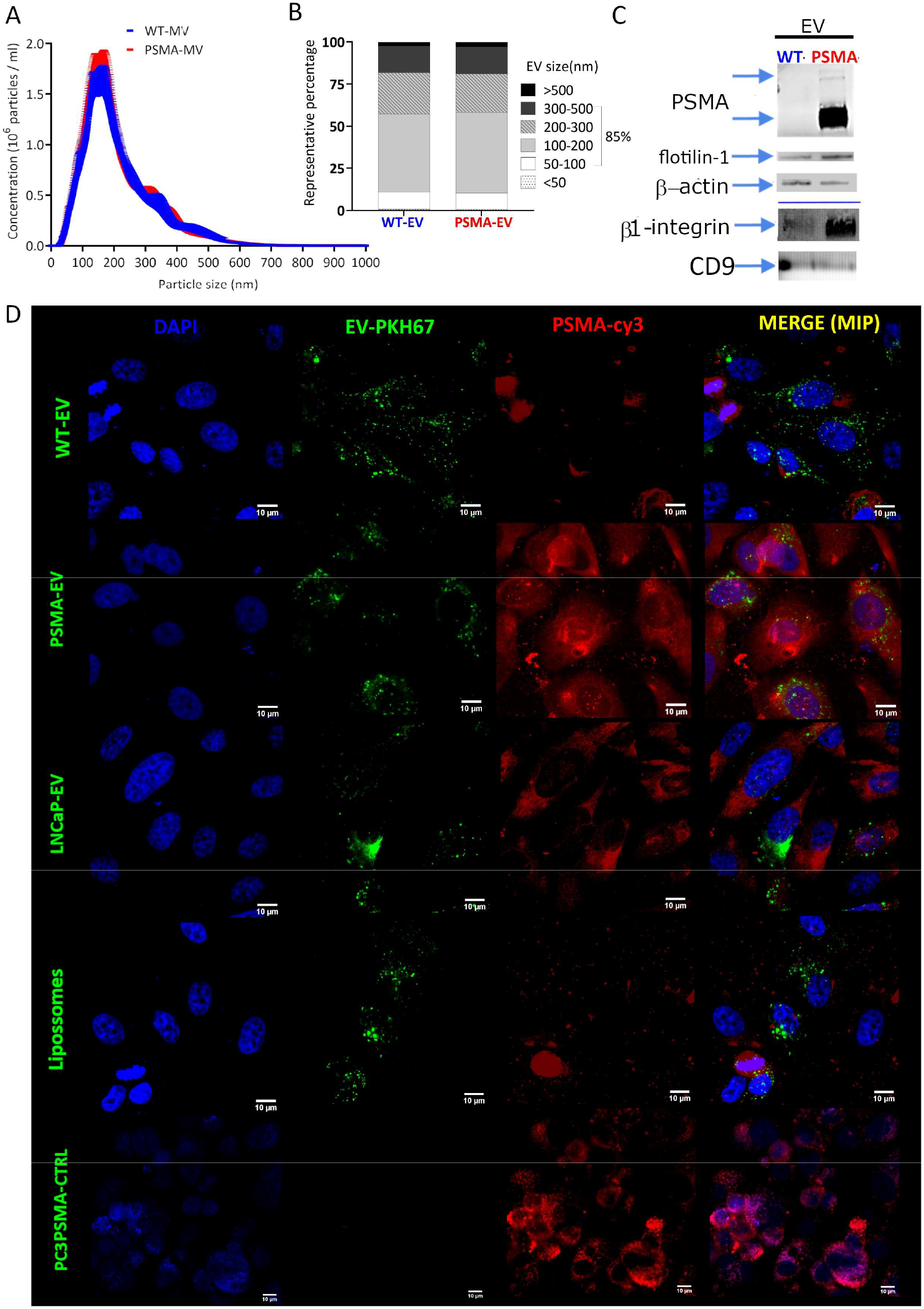
Characterization of EV by size, PSMA expression, and PSMA-transference to recipient cells. **(A)** WT-EV and PSMA-EV size characterization from different EV extractions (n=12) measured by dynamic light scattering analysis showing a similar size range. **(B)** Immunoblot of total EV protein content, separated by native gel electrophoresis, PSMA up to 24 h of PSMA-EV incubation. EV isolations immunoblot to the markers flotillin-1, CD9-tetraspanin, and beta-actin. **(C)** PSMA detected in both monomer and dimer forms via immunoblotting in total protein extraction from PC3WT cells after exposure to PSMA-EV. **(D)** Representative fluorescence microscopy micrographs of PC3WT cells after a 24 h incubation period with fluorescently labeled EV (membrane-label green, PKH67-labeled) followed by staining with a primary PSMA antibody and secondary Cy3-labeled antibody (red) along with DAPI nuclear staining (blue) compared to PC3-PSMA cells (as positive control). Scale bar = 10 µm.

Having established that only PSMA-EV but not WT-EV could transfer PSMA to recipient cells, and that both PSMA- and WT-EV had no differences among hydrodynamic size or uptake rates *in vitro*, we then investigated the biodistribution and *in vivo* effects of EV in healthy adult nude mice using defined EV doses published previously (Lai et al., 2014) (Supplementary Fig.3A-E), for both WT- & PSMA-EV. Consistent with findings *in vitro*, no signs of toxicity were observed in mice: e.g., no changes in body weight, ulceration, bleeding, or labored breathing were observed for up to 25 days of daily tail vein injections of isolated EV. Moreover, WT-EV and PSMA-EV distributed in a similar pattern in all organs analyzed (Supplementary Fig.3). These studies validate and establish PSMA-EV as a vehicle for PSMA transfer to recipient TME cells both *in vitro* and *in vivo*.

### PSMA-EV modify cell types present in the TME to increase secretion of pro-angiogenic factors

Previous studies have demonstrated expression of PSMA in tumor neovasculature (Wernicke et al., 2014) and retinal revascularization models (Grant et al., 2012), suggesting a pro-angiogenic function of PSMA. Shapiro et al. showed that PCa tumors in PSMA-knockout mice were less vascularized. The question arose as to whether EV carrying (and transferring) PSMA could contribute directly to the release of vascular factors in the TME. To elucidate the role of EV in tumor angiogenesis, we first investigated whether EV could directly carry pro-angiogenic factors as a protein cargo. Various human pro-angiogenic factors within EV were quantified using an array (Table 1). No significant differences were observed between WT-EV and PSMA-EV among sequential EV batch isolations (Supplementary Fig. 4A), ruling out the vascular factor cargo transport hypothesis as the cause of the increased pro-angiogenic milieu induced by PSMA-EV. EV mRNA content, evaluated by qPCR, demonstrated an increased amount of PSMA mRNA in EV originating from PC3PSMA cells. That increase in PSMA directly reflected the increased intracellular PSMA transcription levels in the cells and their shed EV (Supplementary Fig. 4B). Elevated mRNA levels detected inside PSMA-EV showed that EV can, in addition to protein, also transfer mRNA to recipient cells. mRNA transfer could directly result in an increase in PSMA translation and expression, potentially modifying the TME, a phenomenon previously demonstrated in exosome EV (Yang et al., 2020).

**Table 1:** Summary of pro-angiogenic factors analyzed through quantibody protein array.

| Factor | Abbreviation |
| --- | --- |
| Angiogenin | - |
| Vascular Endothelial Growth Factor A | **VEGF-A** |
| Angiopoietin-2 | **ANG-2** |
| Epidermal Growth Factor | **EGF** |
| basic Fibroblast Growth Factor | **bFGF** |
| Heparin Binding EGF-like growth factor | **HB-EGF** |
| Hepatocyte Growth Factor | **HGF** |
| Leptin | - |
| Platelet derived Growth Factor BB | **PDGF-BB** |
| Placental Growth Factor | **PIGF** |

A cytokine array was then used to investigate whether incubation of PCa cells or stromal cells (endothelial or macrophages), which are abundant in the TME, with EV would elicit a biological effect (Fig. 2 and Supplementary Fig. 5A-F). Only EV carrying PSMA significantly induced the secretion of VEGF-A and angiogenin in tumor cells. (Fig. 2A, Supplementary Fig. 5A) while additional pro-angiogenic factors secreted by tumor cells showed no significant difference between exposure to various EVs (Supplementary Fig. 5D). Endothelial cells exposed to PSMA-carrying EV showed increased expression of VEGF-A and angiogenin (Fig. 2B, Supplementary Fig. 5B) compared to cells incubated with WT-EV, but no increase was observed in the other eight tested factors (Supplementary Fig. 5D). Macrophages had a distinctive response to both WT- and PSMA-EV exposures, causing equivalent increases in VEGF and angiogenin secretion (Supplementary Fig. 5C). However, only PSMA-EV induced upregulation of the pro-angiogenic factors ANG-2, HGF, and PlGF from macrophages (Fig. 2C and Supplementary Fig. 5C). Placental Growth Factor (PlGF) but not VEGF-A (Adini et al., 2002) has been shown to promote endothelial cell survival. Furthermore, PlGF and VEGF-A are also involved in the polarization of TAM to a tumor-promoting, immunosuppressive M2 phenotype (Acedo et al., 2013).

**Figure 2:**
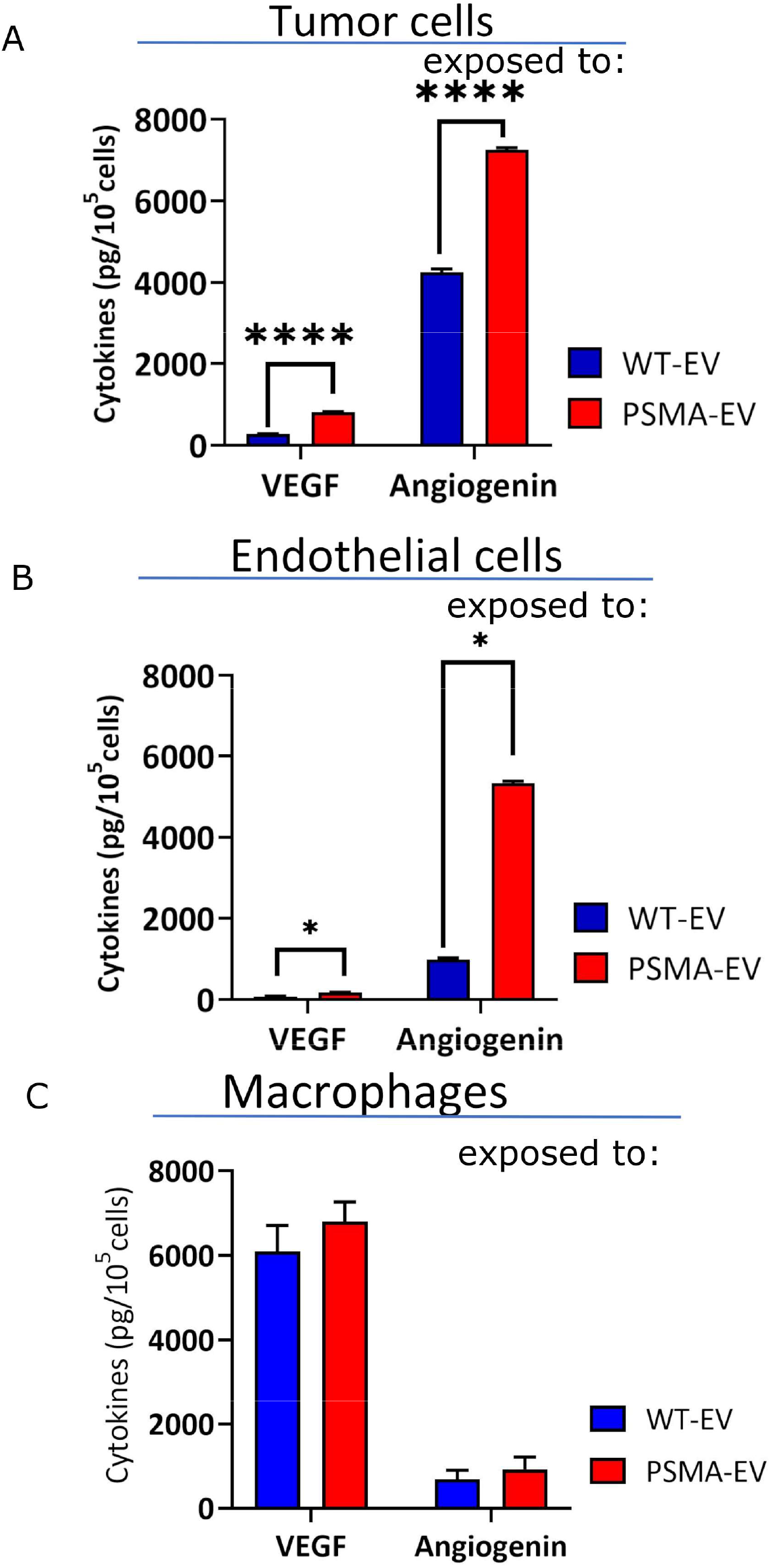
VEGF-A and angiogenin secretion into cellular media after recipient cells exposure to EV. In vitro exposure of (A) tumor parenchyma (PC3WT) or (B) Endothelial cells and (C) macrophages (THP-1) with EV for up to 24 hours measured by protein array. Graphs show the average concentration of VEGF and Angiogenin. N=3 independent experiments for each (2-way ANOVA was used to compare all data, with WT-EV acting as a control; ****P <0.0001 and *P=0.0472).

Demonstrating the role of PSMA expression in regulating VEGF-A, knockdown of PSMA in PSMA-expressing PCa cells (LNCaP) reduced their VEGF-A secretion (Supplementary Fig. 5H). PC3 cells genetically engineered to overexpress human PSMA (PC3PSMA) or CT26 expressing mouse PSMA (CT26mPSMA) exhibited increased VEGF-A secretion (Supplementary Fig. 5I and 5J; respectively). Since pro-inflammatory cytokines can also induce pro-angiogenic activity, we evaluated levels of cytokines involved in macrophage migration, proliferation, chemoattraction, or activation (Supplementary Fig. 5L and 5M). However, our experiments demonstrated no statistically significant differences in the secretion of pro-inflammatory cytokines by recipient cells exposed to WT-EV, PSMA-EV, or liposomes. Our data show that PSMA-EV increase pro-angiogenic signals in stromal cells prevalent in the TME (Supplementary Fig. 5D and 5G), predominantly via increased expression of angiogenin and VEGF-A. These results show that PSMA-EV can modify key cellular mediators of angiogenesis to create a pro-angiogenic milieu in the TME.

### PSMA-EV increased PSMA expression in xenografts

Following the observation of increased pro-angiogenic factors in cells abundant in the TME after exposure to PSMA-EV *in vitro*, we next subcutaneously xenografted mice with PC3WT tumors to investigate the effects of PSMA-EV *in vivo* on primary tumors (Smyth et al., 2015). Once tumors had reached a volume of 0.2 mm^3^, mice received a daily tail vein injection of either WT-EV or PSMA-EV (2×10^10^ vesicles/injection/day/mouse, ten times less than the amount found in the serum of PCa patients) (Padda et al., 2019).

Tumors in mice injected daily with WT-EV or PSMA-EV groups showed similar EV distribution to both tumors and organs, assessed by fluorescence imaging (Fig. 3A), corroborated by *ex vivo* imaging (Supplementary Fig. 6A). PSMA-EV injections increased the level of PSMA up to 4.6-fold in tumors compared to WT-EV. PSMA expression levels were detected using the PSMA-targeting fluorescent agent, YC27-Cy5.5 (Chen et al., 2009) (Fig. 3B). YC27-Cy5.5 showed no significant difference in organ accumulation between tumor-bearing and healthy mice (Supplementary Fig. 6B). *Ex vivo* tumor images in the PSMA-EV exposed group suggested increased PSMA levels (increase in YC-27-Cy5.5 fluorescence, Supplementary Fig. 6B, insert). Fluorescence levels from blood background were subtracted from all analyzed images to remove residual fluorescence of unbound labels (Supplementary Fig. 6C). Fluorescence microscopy (Fig. 3C) confirmed PSMA-positive areas in both tumors and the spleens (Fig. 3D) of the PSMA-EV exposed group. PSMA-positive areas in tumors had a 6.3-fold fluorescence increase when compared to WT-EV. These results are in line with previous reports showing EV accumulation in the spleen of mice bearing PC3WT tumors (Smyth et al., 2015).

**Figure 3:**
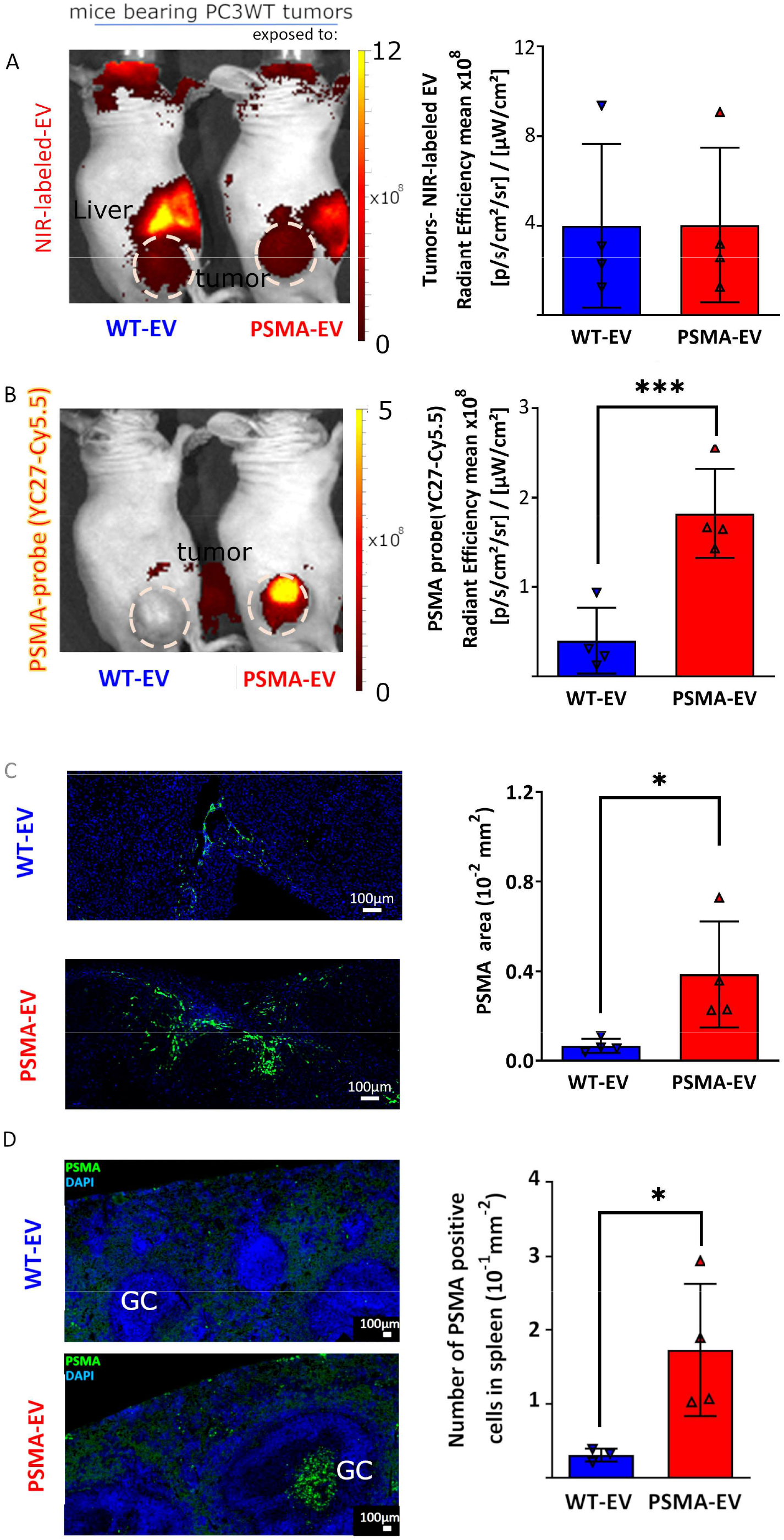
EV distribution in tumors from mice receiving either PSMA-EV or WT-EV. Nude mice bearing PSMA-negative PC3 tumors received daily intravenous injections (tail vein) of WT-EV or PSMA-EV for 25 days. Multispectral fluorescent imaging (MFI) acquisition followed by automated spectral unmixing images are shown to highlight each specific fluorophore-probe. Representative optical images (A-B) BODIPY-CER-EV tumors distribution and quantitation. PSMA (YC27-Cy5.5) fluorescent signal and image quantification. Immunofluorescence microscopy of sectioned tumor (C) and spleen tissue (D), PSMA-positive areas shown by immunofluorescence (green). (The results are representative from the analysis of 20 different fields (20x magnification) and quantification from 2 independent experiments. Graphs represent the average and SD of n = 4 animals each. The unpaired t-test was used to test for significance where *P≤0.04and ***P≤0.004).

### PSMA-EV tumors recruit blood vessels and secrete pro-angiogenic factors *in vivo*

We next explored PSMA-EV effects on recruitment and morphology of tumor vessels *in vivo* by comparing tumor vasculature prior to EV injection (day -1) and at the 9 and 25 day time points in a treatment course of daily EV injections. Vessel recruitment and morphology was assessed non-invasively using raster-scan optoacoustic mesoscopy (RSOM) (Haedicke et al., 2020; Omar et al., 2013). RSOM uses pulsed laser light to induce thermoelastic expansion of hemoglobin molecules, resulting in detectable ultrasound emissions that allow longitudinal noninvasive visualization of microvessel architecture in high spatial resolution as previously described (Haedicke et al., 2020).

On day 25, the density of smaller vessels (indicated by the green color-coded vessels in Fig. 4A) and the vascular branching ratio (Fig. 4B) increased significantly only in mice receiving PSMA-EV compared to tumors before daily injections with EV. Mice were sacrificed after RSOM imaging on day 25, and tumors were excised for 3D microscopy, histology, and protein quantification. Tumor tissue was cleared (Nojima et al., 2017) and stained for the endothelial marker CD31. The whole tissue was then imaged via 3D confocal microscopy. Tissue analysis confirmed an increased number of CD31 positive vessels within the tumors in mice exposed to PSMA-EV compared to WT-EV (Supplementary Fig. 6D). These results confirm that PSMA-EV but not WT-EV induced increased PSMA levels not only in tumors, but also in the vasculature.

**Figure 4:**
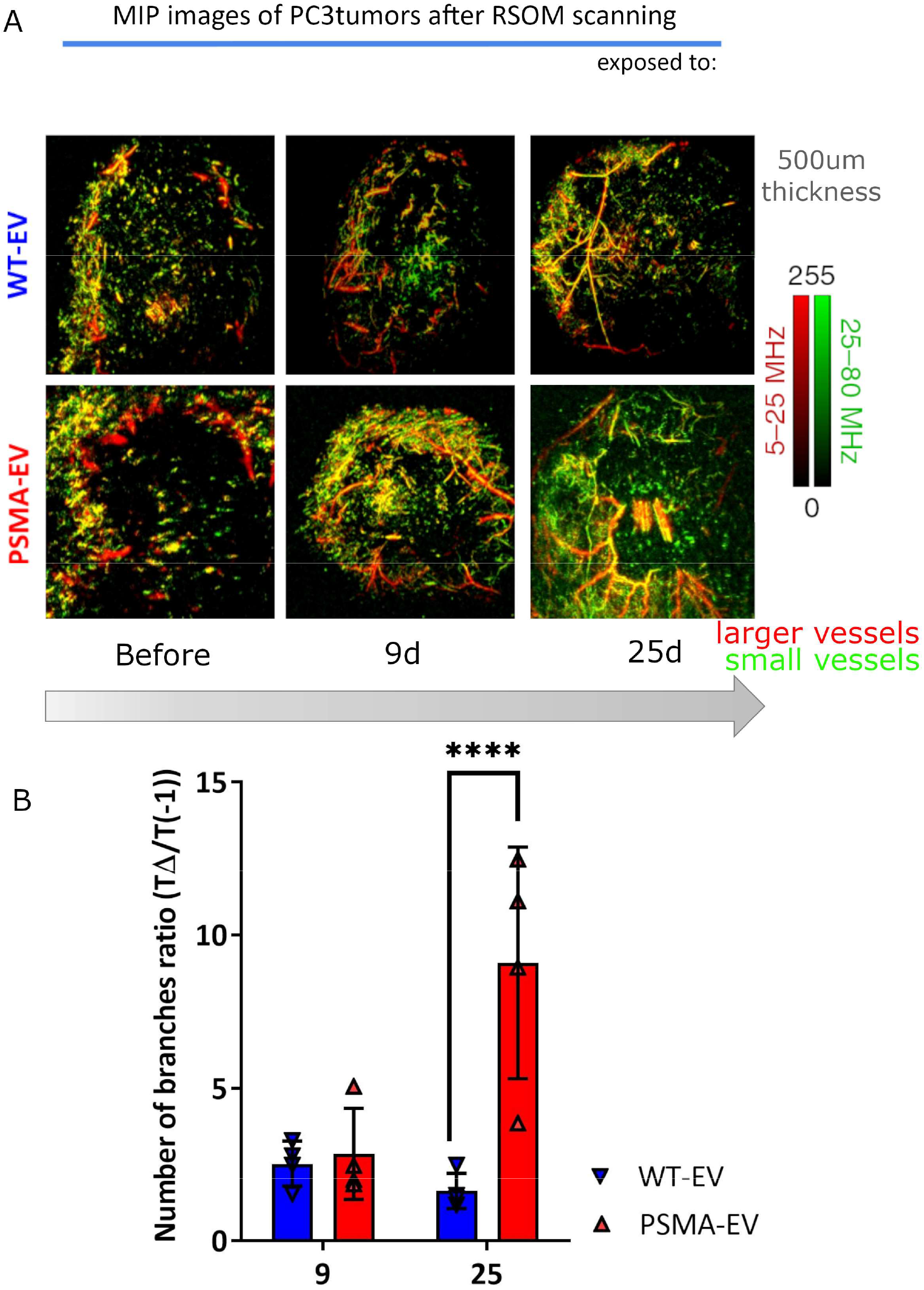
Peripheral and intratumoral microvascular density revealed by RSOM. **(A)** Green and red color-coded vessels formations inside PC3WT tumor xenografts in vivo during EV stimuli. Note visual tumoral neoangiogenesis morphologies, such as glomeruloid microvascular proliferation (GMP) and vascular malformations (VM). **(B)** The representative graphic shows the number of branches inside the tumors at 9 and 25 days after the first intravenous injections to the initial number of branches (*P≤0.002, ****P=0.00001; 2-way ANOVA with Bonferroni correction). Average of three independent experiments.

After confirmation of increased PSMA levels in PSMA-EV-treated animals, we quantified VEGF-A expression in tumor-bearing animals exposed to EV. VEGF-A expression was assessed using a fluorescently labeled VEGF-A antibody (anti-VEGF-A-Cy7). Our work revealed significantly higher VEGF-A levels in animals that received PSMA-EV compared to WT-EV. The fluorescent signal in tumors exposed to PSMA-EV (Fig. 5A-C) was higher than in those that were exposed to WT-EV at 9 and 25 days (1.7-, 2.3-fold respectively; normalized to the fluorescence before the first EV intravenous injection), corroborated by *ex vivo* tumor analysis (Fig. 5C). The overall Cy7 epifluorescent quantitation in organs from both groups in mice bearing tumors or healthy animals showed no uptake differences (Supplementary Fig. 7A-D) showing that the VEGF-A changes are centered in tumor and spleen.

**Figure 5:**
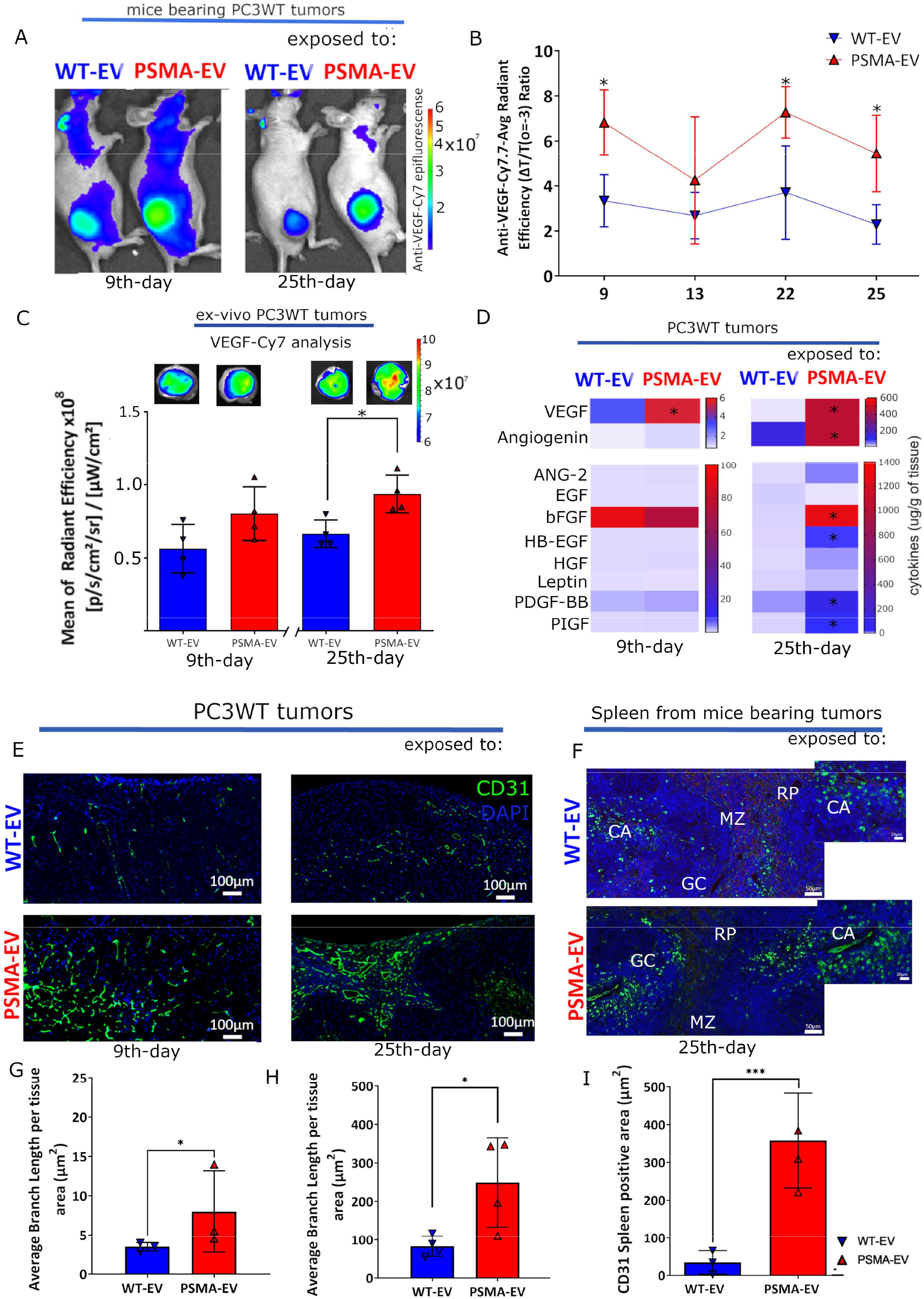
Intratumoral assessment of VEGF-A, pro-angiogenic factors, and microvascular branch length in PC3WT tumor xenografts and spleen in mice receiving daily injections of WT-EV or PSMA-EV. **(A)** Representative in vivo epifluorescence imaging of PC3WT tumor-bearing mice injected daily with EV and injected with **(B)** Cy7-labeled anti-VEGF-A antibody, observed in tumors (24-hours and 96-hours (*) after antibody biodistribution). The images and the quantification are representative of 2 independent experiments (n = 4 mice). All statistical analysis was conducted using 2-way ANOVA (*P≤0.05)) **(C)** Ex-vivo tumor analysis of VEGF-A-Cy7 fluorescent signals (*P≤0.01; unpaired test t) **(D)** A proteomic array assay of 10 pro-angiogenic factors was performed to quantify pro-angiogenic protein levels in tumors. Whole proteins extracts were normalized for respective tumor weight. (*P≤0.01) Each color-code in the heatmap columns represents the average of 4 animals at either 9 or 25 days. (**E**) Representative CD31 immunostaining of tumor tissues at day 9 (left) and day 25 (right) along with **(F)** Representative spleen microvasculature images from mice exposed to EV. (**G & H**) Quantification of vasculature branch length per area (µm^2^) at days 9 & 25 for WT-& PSMA-EV exposed PC3T tumors. (**I**) Quantification of spleen CD31 expression levels from PC3T xenografted mice exposed to WT- or PSMA-EV. Images and the quantification are representative of 2 independent experiments (n = 4 mice, *P≤0.01; ***P=0.001). All statistical analysis was conducted via an unpaired t-test. GC, germinal center; MZ, marginal zone; CA, central artery and RP, red pulp.

A study of extracted protein showed a significant increase (2-fold) in VEGF-A in tumors from the PSMA-EV at day 9 after the EV injection, which increased at day 25 to 100-fold more VEGF-A and angiogenin compared to the WT-EV group. In addition, a significant increase in 6 out of 10 evaluated pro-angiogenic factors was observed in tumors from the PSMA-EV group (Fig. 5D, right column). This upregulation of pro-angiogenic factors in mice receiving PSMA-EV injections was accompanied by changes in the microvasculature architecture in the tumors (Fig. 5E) and also in the spleen (Fig. 5F). There is notably increased microvasculature branching in tumors after 9 and 25 days of PSMA-EV injections (Fig. 5G and H) corroborated by CD31^+^ analysis.

In comparison to WT-EV, exposure of mice to PSMA-EV (up to 25 days) induced morphological vessel changes in both tumors and spleens. We have shown that biochemical changes in tumors are driven by PSMA-EV, including increased levels of PSMA and VEGF-A. In addition, PSMA-EV increased tumoral expression of proteins that have a well-characterized relationship with VEGF-A (bFGF, HB-EGF, PDGF-BB, and PlGF) (Barrientos et al., 2008), which are also increased in acute wound healing and can act as pro-tumoral factors (Foster et al., 2018). Taken together, these biochemical and morphological changes suggest a physiological transition to a more highly vascularized tumor via PSMA-driven alterations to the TME.

To complement and further expand our findings in PCa xenografts, we explored data from The Cancer Genome Atlas (TCGA). The chosen data collection comprised of 492 clinical samples allowing us to compare mRNA levels from PSMA-high versus PSMA-low expression tumors (Hupe et al., 2018). These gene expression TCGA profiles showed a pro-angiogenic molecular signature concomitantly upregulated in PSMA-high samples, similar to with our *in vitro* and *in vivo* results (Supplementary Fig. 8A-F), i.e., with an increased co-expression of VEGF-A, PlGF, PDGF-BB, and ANG-2 (Supplementary Fig. 8A-D). This analysis corroborates a positive and significant correlation between pro-angiogenic gene signatures and increased PSMA-expression in PCa patients. Moreover, PCa patients from this cohort with elevated PSMA, VEGF-A, PlGF, PDGF-BB, ANG-2 (or ANGPT2) gene expressions showed unfavorable overall survival rates when compared to patients with lower genetic expression of the same markers (Supplementary Fig. 8F). As previously observed, PSMA is detected in the tumor neovasculature (Nomura et al., 2014) of non-PSMA expressing tumors; however, its new role in promoting a tumor-supporting environment *in vivo* is a novel and previously unknown aspect of PSMA’s biological function.

Our results demonstrate that repeated application of PSMA-EV resulted in increased VEGF-A and angiogenin levels in tumors and elicit secondary pro-angiogenic factors *in vivo* models. Furthermore, human patient expression data corroborated that increases in PSMA expression correlate to a particular pro-angiogenic molecular signature with diminished overall survival.

Macrophages are known to migrate into tumors and tissues highly expressing VEGF-A (Lewis et al., 2000). The M2-phenotype of macrophages is a source of PDGF-BB in wound healing (Corliss et al., 2016; van der Kroef et al., 2020) but also in the TME (Delprat and Michiels, 2021). CD68 is used as a generic marker for tumor-associated macrophages (TAM) in PCa (Shimura et al., 2000), frequently associated with tumor progression and poor prognosis (Linde et al., 2018). Murine macrophages release EV enriched in nerve growth factor receptor, which are spread through the lymphatics and taken up by lymphatic endothelial cells, ultimately reinforcing lymph node metastasis (García-Silva et al., 2021). This finding hints at a constant communication from tumor cells to cells of the TME, like macrophages and endothelial cells, to promote and support angiogenesis and lymphangiogenesis (Delprat and Michiels, 2021) via induced VEGF-A secretion (Corliss et al., 2016). We aimed to further investigate if PSMA-EV and VEGF-A could prime the PCa TME in a similar fashion as described for melanoma (García-Silva et al., 2021).

Tumors and spleens from the PSMA-EV-group harbored around 2.5-fold more CD68^+^ cells than in the WT-EV group after 25 days of EV injections (Fig. 6A and B). The increased number of CD68^+^ cells coincided with an increased fluorescence from fluorescently labeled anti-VEGF antibodies as well as increased VEGF-A protein levels. Liver and lungs tissues showed no differences in number of CD68^+^ cells (Supplementary Fig. 9A).

**Figure 6:**
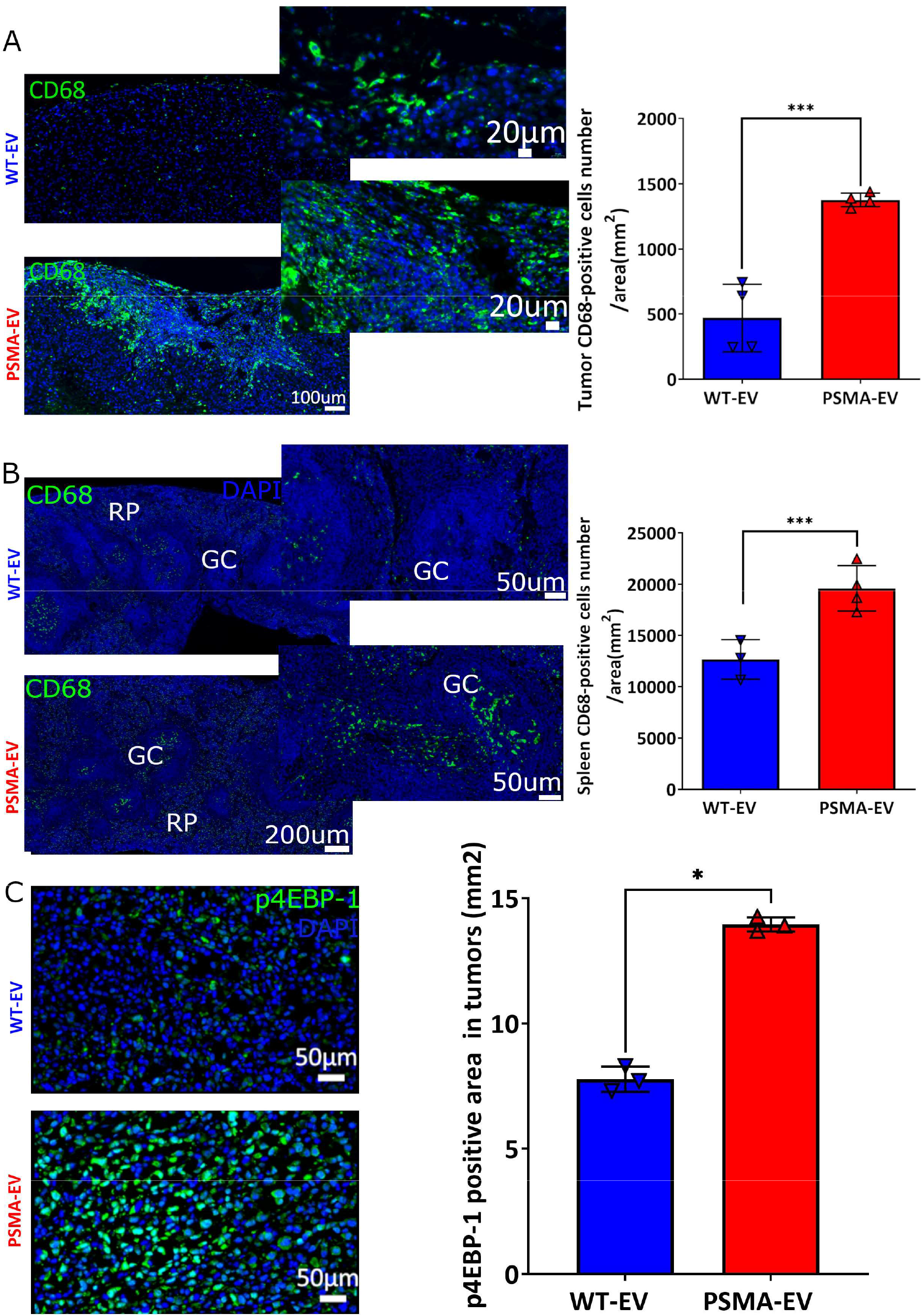
Quantification of CD68^+^ and p4EBP-1^+^ cell levels inside tissues at day 25 post EV exposure. Immunofluorescence images (left columns) and quantification (right columns, graphs) of tissues after daily EV injections into mice showing tumor (A) and spleen (B) CD68^+^ cell recruitment. GC=germinal center, RP=red pulp. (C) p4EBP-1^+^ cytoplasmic and nucleus staining in tumor cells. The images represent tumors from each group ranging from 2× to 40× magnification; quantification results represent repeated independent experiments (n = 3). The data represents the average and SD of animals each. All statistical analysis was conducted by using Student’s unpaired t-test, and the results were considered significant where *P ≤0.05, ***P≤0.001).

Furthermore, tumors from the PSMA-EV group demonstrated increased expression of PDPN (2-fold) and PDGF receptor β (PDGFRβ), multifunctional markers associated with tumor angiogenesis, lymphangiogenesis and LN metastasis (Rolny et al., 2006; Schoppmann et al., 2002; Ugorski et al., 2016) (Supplementary Fig. 9B and D) at day 25. PDGFRβ is also the receptor for PDGF-BB, another pro-angiogenic factor that we detected in protein extracts from tumors exposed to PSMA-EV (Fig. 5D). These findings show that PSMA-EV not only induce and increase tumor angiogenesis by VEGF-A in a PCa model, but additionally enhance recruitment of macrophages to the TME. These PSMA-EV driven TME alterations could further promote angiogenesis and lymphangiogenesis, and potentially facilitate tumor metastasis.

Since we have previously shown that 4EBP-1 phosphorylation from AKT signaling is increased in human PCa tissue samples with higher PSMA expression (Kaittanis et al., 2018), we investigated if PSMA-EV could have the same effect. To this end, we evaluated the phosphorylation status of 18 proteins associated with the AKT signaling pathway. Tumors exposed to PSMA-EV demonstrated significant upregulation of phosphorylated 4EBP-1, GSK3b, RSK1, and RSK2 25 days after daily PSMA-EV injections (Supplementary Fig. 10A and B). The results from the PSMA-EV group confirmed that 4EBP-1 phosphorylation is increased in tumors compared to the WT-EV group (Fig. 6C).

We also questioned if PSMA-EV could result in tumor proliferation stimuli by the MAP kinase (Caromile et al., 2017) signaling pathway. In our mouse model, only after 25 days, differences between groups to the p38 mitogen-activated protein kinase phosphorylation (phospho-p38-MAPK) became apparent. PSMA-EV injected group (Supplementary Fig. 10C and D) have more pp38 in protein tumor extracts than WT-EV tumors that received WT-EV without differences in tumor volume yet (Supplementary Fig. 10E).

Our results show that PSMA-EV alters the TME by increasing pro-angiogenic and -lymphatic factors, driven by the secretion of VEGF-A and angiogenin. These TME modifications are enhanced and altered by time to prime tumor tissue to lymphangiogenesis, that it is sustained by the activation of 4EBP-1 via phosphorylation. Importantly, VEGF can also lead to activation of this pathway (Abid et al., 2004; Zhou et al., 2021).

## Discussion

In recent years, EV have emerged as a new means of PCa detection (Biggs et al., 2016; Minciacchi et al., 2017). However, few studies have explored their pathophysiology. EV have been described as ‘load carriers’ between cells, delivering proteins and mRNA to the TME and modulating signals (Song et al., 2018). PSMA expression and angiogenesis have appeared correlated in experimental and clinical studies, but the significance of this has not been well studied. Our previous work described PSMA as an upstream initiator of the AKT pathway in PCa with an increase in 4EBP-1-phosphorylation (Kaittanis et al., 2018). Here, we investigated whether EV derived from PSMA-positive PCa cells could effectively transfer functional PSMA to primary tumors and stromal cells resulting in re-programming of the TME.

Functional transfer of PSMA from expressing cells to non-expressing cells appears possible. We demonstrated that the EV isolated from PSMA overexpressing PCa cells can deliver PSMA to recipient cells, inducing increased expression of VEGF-A and angiogenin in tumor cells and other cells of the TME, both *in vitro* and *in vivo*. Transfer of PSMA by EV, induces other cellular changes consistent with a PSMA^+^ phenotype. We have previously demonstrated that PSMA induces activation of an AKT signaling cascade via its enzymatic function as a carboxypeptidase, releasing glutamate to trigger metabotropic glutamate receptors with a G protein-coupled receptor function (Kaittanis et al., 2018). The PSMA from EV was integrated without signs of protein degradation and remained able to trigger AKT signaling as described previously (Kaittanis et al., 2018). Since we also detected PSMA mRNA in EV, it is reasonable to suggest that recipient cells also expressed PSMA from the delivered mRNA payload, providing functional PSMA by two different mechanisms: directly, as intact protein by membrane fusion; and as a source of PSMA mRNA for translation by the recipient cell. The EV-mediated conferral of a pro-angiogenic phenotype could be an important biological function of PSMA as a promoter of angiogenesis.

Sustained exposure of tumors to PMSA-EV triggered changes in tumor vascular density and structure, as shown via high-resolution optoacoustic imaging. Unlike intravital microscopy, RSOM does not require an invasive window chamber, allowing undisturbed and non-inflamed transdermal observations of cancer over time (Haedicke et al., 2020; Omar et al., 2013) and can produce higher-resolution images of perfused vessels inside tumors than other non-invasive techniques such as micro-CT and micro-MRI blood volume measurement (Ungersma et al., 2010). Tumors in mice that received PSMA-EV exhibited progressive time-dependent changes in tumor vessel formation, including increased endothelial cell branching and vascular malformations, confirmed by microscopic analysis of tumor tissue. *In vivo* epifluorescence images show a ten-fold increase in intratumoral VEGF-A and pro-angiogenic factors from 9 to 25 days in PSMA-EV exposed groups. This suggests a PSMA-mediated priming of the PC3 TME with released pro-angiogenic factors, effected by EV delivery.

Other effects on tumors were observed on the TME with prolonged PSMA-EV treatment. Increased VEGF-A is known to increase tumor infiltration by macrophages (Lewis et al., 2000), which are associated with tumor angiogenesis (Li et al., 2015) and poor patient prognosis (Minami et al., 2018). Tumors from mice exposed to PSMA-EV for 25 days demonstrated abundant CD68^+^ staining of tumor sections, This VEGF-A expression further increased 100-fold at day 25 compared to levels prior to PSMA-EV injection, highlighting the capability of PSMA-EV to shape a microenvironment favorable to angiogenesis and to alter the local immune environment through influx of macrophages (Fig. 7). We also detected increased expression of the PDGF-BB receptor, PDGFRβ, within tumor tissue from mice treated with PSMA-EV. Increased PDGF-BB expression strongly activates pericytes (Gaceb et al., 2018) endothelial cells and Lymphatic Endothelial cells (LEC) in a synergistically mode to induce neovascularization (Cao et al., 2003).

**Figure 7:**
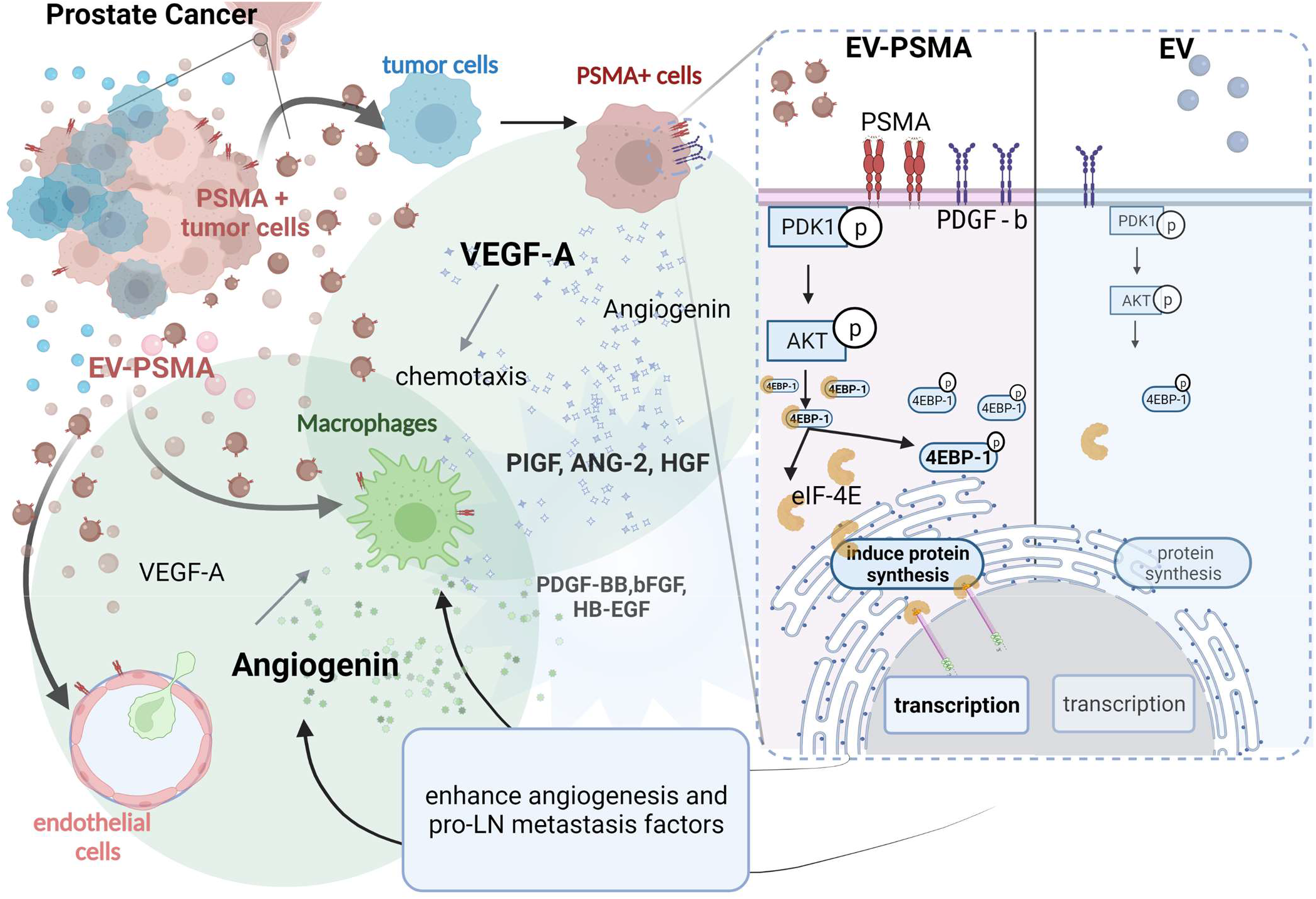
PSMA in EV transfers its biological activity to recipient cells. PSMA-EV activates a switch in the tumor microenvironment that drives prostate cancer angiogenesis at early time points in PCa. This oncogenic switch is activated by increased secretion of pro-angiogenic factors to TME cells ultimately leading to increased 4EBP-1-phosphorylation and then boosting the TME to produce early pro-angiogenic signals (Figure made with Biorender.com software).

Our findings are novel because we show that PSMA-EV can shape the TME, however, our results did not show statistically significant differences in tumor volumes before 25 days, nor did it test for changes in survival. Adding to this, short-term daily exposure to EV (9 days) did not increase phosphorylation of survival factors (such as p38, p53, PTEN, and others), regardless of whether PSMA was present in the EV or if VEGF-A secretion into the TME was increased. Our model shows that PCa produces VEGF and Angiogenin factors modifying TME cells prior promoting uncontrolled tumoral cell proliferation. In contrast to this, the phosphorylation of p38, associated with PSMA expression and an increased cell proliferation status (Huth et al., 2017) was showed before., For further studies a more prolonged PSMA-EV exposure at later time points (more than 25 days) can improve our data. However, VEGF-A secretion can also be driven by hypoxic areas within tumors, however, the model used here did not reach typical hypoxic sizes and was outside the scope of this investigation. PSMA-EV alone cannot be used to determine prognosis in PCa as it was already showed (Krishn et al., 2019; Padda et al., 2019).

There is a novel implication from our results based on the fact that PSMA-EV triggered VEGF-A is also implicated in lymphangiogenesis (Hirakawa et al., 2005) to facilitate tumor spreading to the lymph nodes (Hirakawa et al., 2005). LN metastasis is an important indicator of prognosis (Datta et al., 2010; Jansen et al., 2021).The normal prostate epithelium has relatively few lymphocyte within the tissue, but this increases 100-fold(Dikov et al., 2015) in inflammatory prostatitis, a preconditioning to PCa and further changes to LN activation to predict negative outcomes (Kamel et al., 2020) and staging (Jansen et al., 2021). A recent paper by García-Silva et al.(García-Silva et al., 2021) demonstrated that EV derived from melanomas are delivering nerve growth factor receptor (NGFR) to promote LN metastasis. The authors showed that acutely exposed melanomas models to melanoma-derived-EV elicited Lyve-1 (a dual EC and LEC marker), PDGF-BB, and PDGFRβ increases, resulting in metastasis. Interestingly, in our study PSMA-EV increased ^18^F-FDG uptake in LN of mice after daily injections up to 25 days. In addition to this, increased presence of macrophages as well as PDPN, PDGF-BB, and PDGFRβ, can all promote neovascularization. These factors also contributed to activation of pericytes and have been shown to activate also lymphangiogenesis (Cao et al., 2004; Datta et al., 2010; García-Silva et al., 2021; Kuroda et al., 2008). In patients, lymphatic vessels are frequently observed around tumor tissues coinciding with areas of neoangiogenesis and could possibly be the starting point for PCa lymphatic metastasis (Hirakawa et al., 2005; Lilis et al., 2018; Zeng et al., 2005). The PSMA protein (and PSMA mRNA) is delivered via EV to result in AKT-activation, i.e., 4EBP-1-phosphorylation mainly in tumor cells, to enhance microvessel density (De Benedetti and Graff, 2004) and enhance secretion of pro-angiogenic, and pro-lymphangiogenic factors. The AKT activation followed by 4EBP-1-phosphorylation in PCa is correlated with tissue invasiveness and lymph node metastasis (Brakenhielm et al., 2007; Chen et al., 2015; Graff et al., 2009).

In conclusion, we show that within a timeframe of 25 days in mice, PSMA-EV derived from PSMA-positive tumor cells induced pro-angiogenic and pro-lymphangiogenic factors in PCa by transferring PSMA and conferring a PSMA-positive phenotype on both tumor and stromal cells. As a result, enhanced vessel recruitment and angiogenic factors are sustained by increased cell recruitment to the TME and 4EBP-1-phosphorylation further promote ongoing translation. This work bridges a gap from previous studies to show that the TME is modulated by PSMA-EV beneficial to PCa around a 25-days window in already established tumors by far exceeding the observation time frame of prior studies in vivo.

## Materials and Methods

### Cell culturing for mice models and EV analysis

As PSMA-EV donors, we used PC3 cells that were retrovirally transfected to express human PSMA (named here as PC3PSMA)(Kaittanis et al., 2018) or LNCAP and 22RV1 purchased from ATCC. For donors of non-PSMA expressing (WT-EV), we used PC3WT, the non-PSMA PCa. We also used the PSMA-mirRNA-inhibited LNCaP cells from our previous work(Kaittanis et al., 2018) to measure VEGF-A secretion compared to LNCaP cells as further described. PCa cells were cultivated in F-12K or RPMI, pH 7.4, supplemented with 10% heat-inactivated fetal bovine serum (FBS) (Gibco, Life Technologies, Carlsbad, CA) with 2 µg mL^-1^ of puromycin. RAW264.7 cells were cultivated in DMEM high-glucose media, while THP-1 cells were grown in RPMI, pH 7.4, with 10% heat-inactivated FBS as a supplement (Gibco, Life Technologies, Carlsbad, CA) and 1% penicillin and streptomycin (Gibco, Life Technologies, Carlsbad, CA). Human umbilical vein endothelial cells (HUVEC and HUVEC-RFP) were cultured in vascular EnGS medium (Vasculife, Lifeline Cell Technology, CA) supplemented with 2% low heat-inactivated FBS. We also used the colon carcinoma cell lineage CT26 or retrovirally transfected to express murine PSMA (CT26mPSMA) in RPMI, supplemented with 10% of FBS. All cells were kept in a 95% humidified atmosphere with 5% CO2 at 37°C. For confocal experiments with EV and cells, PC3 cells were plated over Nunc® Lab-Tek® chamber slides to 70% cell density on the day of EV incubation. For PC3WT cells, for inducible pro-angiogenic factors experiment, we treated the cells with 0.02µg of human recombinant PSMA (Recombinant Human PSMA/FOLH1 Protein, #V 4234-ZN-010, R&D, US) up to 72h followed by EV isolation and protein extraction. A mycoplasma routine quality test was performed once per month for each cell line by collecting 1 mL of media, centrifuging at 800 x g for 10 min to remove cells or apoptotic bodies, and applying the MycoAlert™ detection kit (Lonza) according to the manufacturer’s instructions.

### EV isolation by differential centrifugation and size characterization

The respective human prostatic cell lines were deprived of FBS to collect EV for 3 to 5 days, as described above. The method chosen was inspired by the literature(Kowal et al., 2016), and a scheme with a summary can be seen in Supplementary Fig.1A. The media was collected in tubes and apoptotic bodies were removed by two serial centrifugations at 800 xg for 15 min each, at 4°C. Afterward, samples were submitted to centrifugation at 11,600xg at 4°C for 120 min followed by 2 washing steps (11,600 xg at 4°C for 60 min), and resuspension in filtered PBS and stocking at −80°C. The size characterization and polydispersion index were measured using a Zetasizer Nano series (Malvern Inc.), with 15 runs of 10 s each for every sample at 4°C. EV quantification and characterization were done by nanoparticle tracking analysis (NTA) at 21°C using a Nanosight NS300 equipped with a 488-nm laser and a sCMOS camera to every isolation/batch. At least 3,000 events in six different regions were measured to obtain EV concentration. For each EV approach, the average of all counted samples was considered as the amount of EV. For each EV characterization, six videos were recorded.

### Liposome preparation for *in vitro* and *in vivo* uptake studies

Liposomes were constructed of cholesterol from ovine wool, 1,2-dipalmitoyl-sn-glycero-3-phosphocholine (DPPC), and sphingomyelin in a 60:30:10 molar ratio 100 mg total, from Avanti Polar Lipids (Alabaster, AL). Lipids were dissolved in chloroform and rotary evaporated to produce a lipid film, which was rehydrated with 1X DPBS and cup sonicated at 15kJ over 90 s and then extruded through a 0.22 µm filter. Liposomes were characterized by DLS and NTA, as described above.

### Protein extractions and western blotting for protein content analysis *in vitro* and *in vivo*

EV, cell, or tissue samples were homogenized in two types of buffers: one specifically for EV and cells (FNN0011, Thermo Fisher Scientific Inc., New York, NY) and the other one for tissues (FNN0071, Thermo Fisher Scientific Inc., New York, NY) as indicated by the manufacturer for full protein extraction. All lysis buffers contained 2 μg mL-1 aprotinin (#78432, Thermo Fisher Scientific Inc., New York, NY), 1 mM PMSF (329-98-6, Sigma-Aldrich, phenylmethanesulfonylfluoride), and protease and phosphorylase inhibitor pierce tablets (#A32957, Thermo Fisher Scientific Inc., New York, NY). The protein isolation was obtained by centrifugation with 13,000 g at 4 °C for 15 min. Protein concentration was determined using the bicinchoninic acid (#23225, Thermo Fisher Scientific Inc., New York, NY) macro- and micro methods. After extraction and quantification, proteins were used for protein arrays or western blot. Total protein extracts from each sample (20-50 µg) were electrophoretically separated in an SDS-PAGE and electroblotted onto a PVDF membrane according to standard procedures. Membranes were then blocked using the commercial casein blocking buffer (#37593, Thermo Fisher Scientific Inc., New York, NY) or a buffer containing BSA excess followed by several washes in a buffer containing 0.1% of Triton-X and up to 3% of BSA. Further steps included primary antibody incubations (anti-PSMA-AbCam (AB1971,YPSMA-1, 1:2,000), anti-PSMA (J591, 1:5000) or anti-PSMA (2µg/mL, MAB4234, R&D Biosystems); anti-beta-actin-Sigma-Aldrich (AC74, 1:4,000), anti-flotilin-1(#3253), anti-CD9 (1:1000, CBL 162), anti-beta-1-integrin (1:1000, Chemicon international) and p38 and phosphor-p38 from cell signaling (1:2,000); anti-VEGF-AR2(#44-1053G) and anti-phospho-VEGF-AR2 (#44-1052)-Thermo Fisher Scientific Inc. (New York, NY) (1:1,000) in PBS with 0.1% Tween and 1% BSA overnight at 4 °C. Membranes were triple washed with PBS with 0.1% Tween and incubated with specific Licor anti-mouse or anti-rabbit fluorescent secondary antibody (925-32212, 1: 4,000) with 700 nm or 800 nm emission peak. Reactive bands were detected using a Licor Odyssey infrared scanner. The images were quantified using the Licor software with inverted black-and-white information and ImageJ.

### RNA extraction from EV for RT-qPCR reaction

EV were isolated from WT and PSMA-overexpressing PC3 cells as described above. RNA was collected from both EV using the RNA isolation kit from Exiqon following the manufacturer’s instructions. Then, cDNA was synthesized from 1 µg of RNA using High Capacity kit (ThermoFisher Scientific, Waltham, MA) and real-time qPCR was performed using SyBR Green (SYBR Green Master Mix; Thermo Fisher Scientific, Waltham, MA) chemistry for PSMA gene (Forward: 5’-TTG GAA TCT TCC TGG AGG TG-3’; Reverse: 5’-CTG CTA TCT GGT GGT GCT GA-3’). Flotilin-1 was used as endogenous control (Forward 5’-GAA GAC GGA GGC TGA GAT TG-3’; Reverse 5’-ATC CGT GCA TCT TTT TGG AC-3’, respectively). Independent EV isolations (batches) from PC3 WT or PSMA-overexpressing cells were used in each reaction. The relative gene expression (fold change) was obtained by the 2^delta-delta Ct method, according to described in the literature (Livak and Schmittgen, 2001), using the wild type-derived EV as the reference and EV from PSMA-overexpressing cells as the target sample to fold-increase calculations. The reaction was performed in the ABI 7500 Real-Time PCR machine (Thermo Fisher Scientific, Waltham, MA).

### EV or liposome labeling to PKH (67 or 26) and BODIPY-CER-TR for *in vitro* and *in vivo* uptake assessment

Fluorescent EV for flow cytometry, microscopy or *in vivo* studies were obtained by incubating EV with 1 nM of the near-infra-red fluorescent BODIPY-TR-ceramide (Ex580, Em617, C39H50BF2N3O4S, Thermo Fisher Scientific Inc., New York, NY) at 37°C for 30 min. After the incubation, the EV solution was filtered through exosome spin columns (MW 3,000 kDa) (Thermo Fisher Scientific Inc., New York, NY) to wash away the free dye. The same technique was used to conjugate the control liposomes to BODIPY-ceramide. For microscopy studies, EV were labeled with a PKH67 dye labeling kit-green (or PKH26 dye labeling kit-red) following the manufacturer’s instructions (Sigma-Aldrich). Briefly, a small volume of EV was incubated with 0.2 mM of PKH67 in a very diluted ethanol solution, followed by successive centrifugation in spin columns as described above to eliminate the free dye from the preparation. Stained EV were quantified using NTA analysis. PC3WT cells (2×10^4^ cells/well) were cultivated as described above in four-chamber glass slides (Millipore) then incubated with PKH67-labeled EV (2×10^9^) for 24 h followed by extensive washes with PBS before microscopic imaging. Z-stack images were acquired with a step size of 0.9 µm with a Leica TCS SP5 confocal microscope (Leica Microsystems, Inc., Buffalo Grove, IL).

### *In vitro* EV uptake and PSMA transference to cells evaluated by Flow cytometry

After EV were labeled with BODIPY-CER, they were incubated with different cells for 30, 60, or 90 min, followed by washing with PBS, pH 7.4 at 4°C. Then the cells were harvested using trypsin and centrifuged at 150 xg for 10 min. The cell pellets were suspended in supplemented media specific for each cell type. The flow cytometry result was achieved by comparing cells that were not incubated with labeled EV (named as controls) with cells incubated with labeled EV. The anti-human-PSMA-APC antibody-clone REA408 (or Isotype control, recombinant human IgG1-APC from Miltenyi) was used to label PSMA in cells without or with EV incubation. Briefly, we incubated the antibody in a proportion of 1:11 for 10^7^ cells/100 µL as indicated by the manufacturer. Every batch of cells was compared with cells without labeling (negative control) and the isotype control. The data were acquired using a Miltenyi FACS MACSQuant-Analyzer. The data analysis and graphical summarization were achieved by using FCS express software analysis 6.0 (De Novo Software, Glendale, CA).

### Negative-PSMA Tumor mouse models and daily EV injection to mimic tumor EV-shedding

All procedures were conducted by the ethical principles and institutional guidelines of and approved by the IACUC of Memorial Sloan Kettering Cancer Center as well as the Guide for the Care and Use of Laboratory Animals by the National Institutes of Health (NIH). All animal procedures were performed under anesthesia by inhalation of a 1–4% isoflurane-oxygen mixture (Baxter Healthcare, Deerfield, IL). The EV biodistribution in healthy mice was performed in Six-week-old male Balb/c nude outbreed mice (n=4 each group) using BODIPY-CER-EV injected daily through tail vein, and they were followed by images as described below. The healthy mice were submitted to PET imaging prior to RSOM and iVIs as described below. Also, six-week-old male Balb/c nude outbreed mice were obtained from Taconic Laboratories for experiments with EV in tumor-bearing mice. These mice received a subcutaneous inoculation of 2×10^5^ PC3WT cells in the lower right flank, and tumor growth was determined by measuring the length and depth of the tumor with a digital caliper. In these mice, after 33 days, all mice that developed measurable tumors were imaged for a baseline and time points value by RSOM or IVIS as described below. The mice were designated to specific groups and injected daily as described by other authors (Bruno et al., 2009). Two groups were created with daily injections of PSMA-EV and the other WT-EV (2×10^10^ vesicles/injection/day/mouse, 10 times less than the amount quantified in serum of PCa patients according to literature) (Padda et al., 2019). Tumor volume was calculated based on an approximation of an ellipse volume. At the end time point, mice were sacrificed in constant CO2 air flow for 15 minutes and tissue samples collected for further analysis.

### LN Imaging

Positron emission tomography and computed tomography (PET/CT) images was carried using The Siemens Inveon®. Briefly, nude mice under anesthetic conditions received a retro-orbital intravenous injection of ^18^F-fluoro-2-deoxyglucose ([^18^F]-FDG. The tracer was produced by Department of Nuclear Medicine at Memorial Sloan-Kettering Cancer Center (MSKCC) Radiochemistry Core Facility. For both groups 300μCi per mouse (11.1MBq) was injected in all experiments and mice were imaged after 60-min of biodistribution. PET software with decay correction followed by a DICOM and pictures generations by the Amira 4.1 software (FEI Visualization Sciences Group, Bordeaux, Zuse Institute, Berlin, Germany).

### Angiogenesis morphology detection by Raster-scan optoacoustic mesoscopy (RSOM)

RSOM was performed with a scanner from the Institute for Biological and Medical Imaging at the Helmholtz Zentrum in Munich, Germany, as previously described(Haedicke et al., 2020; Omar et al., 2013). In short, the anesthetized mouse was placed onto a mouse bed and into a warm water bath with the head above the water. The tumor area was fastened with a transparent foil to suppress breathing motion. The tissue was then illuminated with a fast nanosecond pulsed laser (2 kHz, pulse energy of 1 mJ) at a wavelength of 532 nm and the generated ultrasound signal was detected using a spherically focused 50 MHz detector with a bandwidth of 5-80 MHz. The field of view was defined based on tumor size, and the scan was performed in a 20 µm step size raster with a maximum penetration depth of the laser of about 2 mm. After data acquisition, the raw signals were divided into two frequency sub-bands of 5-25 MHz and 25-80 MHz for reconstruction to distinguish bigger and smaller vessels within the tissue(Haedicke et al., 2020; Omar et al., 2013) They were subsequently assigned to different colors, with red displaying bigger and green depicting smaller vessels. During the reconstruction process, voxel sizes of 20×20×5 µm^3^ and speed of sound of 1540 m/s were applied. Image processing was done using the Hilbert transform along the depth axis of the image and utilizing a Wiener and Median filter to improve the overall signal-to-noise ratio. The maximum intensity projection images of the tumors were then transferred to ImageJ, the low (colored in red) and high (colored in green) frequencies were merged. All data was quantified for vascular branching using the Fiji tubeness tool(Costantini et al., 2015; Sato et al., 1998) followed by skeleton (3D/2D) analysis.

### *In vivo* EV-BODIPY, YC-27 Cy5.5 and VEGF-A-Cy7 detection by molecular imaging

All fluorescent animal imaging was conducted using an IVIS Spectrum CT small animal imaging system (Xenogen, Alameda, CA). The VEGF-A accumulation in tumors was monitored in animals injected with bevacizumab-Cy7 or Isotype-IgG-Cy7 (10ug/mouse) as a control 24 h before image acquisition. For the antibody labeling, 2 mg (13.3 nmol) of bevacizumab was buffer exchanged with PBS (pH 8.3) using 0.5 ml Amicon filters (30 kDa) at 4°C and 14,000 xg. Bevacizumab or IgG concentration was adjusted to 1 mg/ml, and sulfo-Cy7 NHS ester (16 µL, 5 mM in DMSO) was added. The reaction mixture was shaken in the dark for 1h at RT. Another buffer exchange was performed with PBS (pH 7) to remove the free dye. The purity of the conjugate was confirmed using gel electrophoresis, and the fluorescence emission spectrum was further verified using a SpectraMax M5 spectrophotometer. Animal images were acquired using an ICG filter (710-760 nm excitation, 810-875 nm emission), automatic exposure time, binning of 8, and a wide field of view (25 × 25 cm). The fluorescent images were normalized to photons per second per centimeter squared per steradian (p s^−1^ cm^−2^ sr^−1^). All images were captured as indicated inside parenthesis after distribution of anti-VEGF-A-Cy7 biodistributions at days, −3 (24h), 9 (24h),13(96h),21(24h) and 25(96h) after daily EV injections into tail vein. PSMA-YC-27-probe-Cy5.5 where injected 24h before images acquisition. BODIPY-TR-CER-EV were injected daily as described above. For quantitative purposes, the background fluorescence was individually subtracted for each group, and automated spectral unmixing was applied. The Living Image 3.0 software (Xenogen, Alameda, CA) was used for data quantification.

### Tissue clearance and staining for ex-vivo angiogenesis quantitation by confocal microscopy

To increase the laser penetration and enable the collection of more data from the confocal imaging of thick tissue samples, clearing of the tissues was performed. The clearing of the samples was done at the Molecular Cytology Core Facility, MSKCC, using CUBIC (Susaki et al., 2014) 300 µm thick vibratome sections were fixed in 4% paraformaldehyde and incubated in CUBIC at least for 1 hour shaking at RT (CUBIC solution is a mixture of 5% N,N,N,N’-tetrakis (2-hydroxypropyl)-ethylenediamine, 5% urea and 80% of Triton-X-100 (g/g/v) as described by (Nojima et al., 2017) The samples were blocked with 2 % BSA in PBS (pH 7.4) for 2 h and washed in PBS, followed by incubation with a CD31 (rat monoclonal antibody from Dianova, cat. # DIA-310) for 2 days at 37°C, with rocking. Afterward, the samples were washed 3-5 times at RT for 1 h in PBS containing 0.1% Triton-X-100, followed by incubation with a secondary goat anti-rat Alexa Fluor-568 antibody for one day at 37°C with rocking. The z-stack images of the whole tissue were acquired with a 5 mm step size in a Leica TCS SP5 confocal microscope (Leica Microsystems, Inc., Buffalo Grove, IL). Images were segmented and volumetrically reconstructed with Imaris software (Bitplane, Zurich, Switzerland).

### Immunofluorescence staining for ex-vivo angiogenesis quantitation by fluorescence microscopy in tissue slices

For immunofluorescence staining of tissue slices, tissues were fixed in 3.7% formaldehyde in PBS with a pH of 7.4. After paraffin embedding, they were sliced with 5 mm thickness and mounted onto glass slides. After three washing steps with PBS, the slices were incubated with primary antibodies: rabbit polyclonal anti-PSMA/GCPII (1:150) (M362029-2; DAKO, US), anti-CD68 (1:250) (Ab5212, Abcam, Calsburg, CA), hamster monoclonal anti-podoplanin (1:150; cat. number ab11936 Abcam, Calsburg, CA), rabbit monoclonal anti-PDGFRβ (1:250; cat. number, 3169 Cell Signaling) or rat monoclonal anti-CD31 (1:350) (cat. number DIA-310 Dianova). The detection of PSMA/GCPII, CD68, podoplanin, PDGFRβ, CD31 were performed at Molecular Cytology Core Facility of Memorial Sloan Kettering Cancer Center, using Discovery XT processor (Ventana Medical Systems). After another three washes with PBS, the slices were incubated with a secondary goat-anti-mouse or rabbit Alexa-488 or Alexa 563 antibodies or goat-anti-mouse-HRP (Invitrogen, Life Technologies, Carlsbad, CA) for 2 h followed by DAPI cell nuclei staining or Hematoxylin staining. For each batch, a negative control (isotype for the primary antibody) was included to validate the results. All stained slices were thoroughly scanned and reconstructed with Panoramic Viewer (3D Histotech, Hungary). For CD31 vessel branching analysis, the FIJI tubness(Costantini et al., 2015; Sato et al., 1998) tool was used to quantify vessel branching followed by skeleton (3D/2D) analysis.

### Angiogenesis and AKT/mTOR protein arrays for factors detection

A 50 μg of whole protein lysate or 1 mL of cell culture media was used to determine the pro-angiogenic pathways by the Quantibody human array kits (RayBiotech Inc., Norcross, GA) according to the manufacturer’s instructions. For angiogenic analysis, the Quantibody Angiogenesis array-Q1 or array-Q100 was used. Briefly, the supplied glass slides were blocked with a blocking buffer solution, and then 1 mL of equal amounts of cell culture media was added over the glass slides and incubated overnight at 4°C. The next day, the slides were extensively washed 8 times for 5 minutes each in washing buffer provided by the kit, followed by the detection antibody cocktail. After the second round of washes, the fluorescent dye-conjugate (Cyanine 3) was added. The slides were read and quantified by sending them to Raybiotech Inc., followed by using an analysis tool from the same incorporation. When applicable, the factored amount was corrected by several cells plated. For AKT/mTOR analysis, the whole protein extract of the tumors was collected as described above, and the Human/Mouse AKT Pathway Phosphorylation Array C1 (AAH-AKT-1-2, RayBiotech Inc., Norcross, GA) was used. Briefly, the PVDF membranes were blocked using blocking buffer solution, and 50 µg of protein were applied to each membrane, which was then incubated overnight at 4°C. The next day, the slides were extensively washed 8 times for 5 minutes each in the washing buffer, followed by the detection antibody cocktail. After the second round of washes, the fluorescent dye conjugate (near-infrared LI-COR antibody with 800-nm emission) was added. For quantification, an LI-COR Odyssey infrared scanner was used, followed by mathematical analysis using the equation indicated by the manufacturer. All data were normalized to an internal positive control tumor and compared to mice with daily injections of WT-EV to show the fold-increase.

### ELISA for specific VEGF-A pro-angiogenic detection

Cells were cultivated as described above and plated in well plates for 24h with media without FBS. Before enzyme-linked immunosorbent assay (ELISA) analysis, conditioned media were centrifuged twice at 800 xg for 10 min each to remove floating cells and apoptotic bodies. Equal amounts of conditioned media were measured using a VEGF-A kit (Peprotech, Rock Hill, NJ) that recognizes human VEGF-A (hVEGF-A) or mouse VEGF-A (mVEGF-A) according to the manufacturer’s instructions. All data were determined based on a standard VEGF-A dilution curve and normalized by the number of cells.

### Gene expression and molecular signature analyses

The cancer genome atlas (TCGA) data sets for 550 prostate adenocarcinoma samples (PRAD, primary solid tumor) were downloaded from the Firehose Broad GDAC portal (http://firebrowse.org/). Tumors with PSMA levels more remarkable than the median expression levels (PSMA levels > 8) were defined as “High-PSMA” tumors, whereas the tumors that fell below the median (PSMA levels < 8) were defined as “Low-PSMA” tumors (RSEM normalized RNA-Seq, gene/GAPDH). For standard co-expression analysis was considered Spearman’s correlation coefficient was used for assessing the correlation between two gene expression levels. Statistical analysis and data visualization were carried out using R (version 3.3.2). Disease free survival correlation analysis was also performed from the TCGA data set using gene expression profiling interactive analysis (GEPIA) web server (Tang et al., 2017) and “High-PSMA” vs “Low-PSMA” tumors (n=320) plus VEGFA, PlGF, PDGFBB and ANGPT2 genes signatures identified as 5-signatures group.

### Statistical analysis

All data were individually analyzed by different statistical methods using a non-parametric test, and significance calculated as noted in each figure legend. All analysis was conducted using GraphPad Prism version 8.0 for Windows^®^ (GraphPad^®^ Software, San Diego, CA) and Microsoft Office^®^ Excel software 2007 (Microsoft, Redmond, WA).

## Acknowledgments

This study was supported by FR01CA212379, FAPESP (# 2015/11808-4 and # 17/01130-6), and Capes (# 88881.119225/2016-01) grants. We thank Dr. Pat Zanzonico and Valerie Longo from the MSKCC Small-Animal Imaging Core Facility, which is supported in part by the MSKCC NIH/NCI Cancer Center Support Grant (P30-CA008748). The authors would like to thank Afsar Barlas and Dmitry Yarilin for the immunohistochemistry staining and Dr. Polly Gregor for providing the CT26-mPSMA cells.

## Authors’ contributions

CMLM conducted and analyzed directly or indirectly all experiments and wrote the main manuscript under the supervision of JG. MS helped with the manuscript corrections and immunofluorescence. KH conducted RSOM experiments and helped with the manuscript writing. FPS and TLSA helped to proofread the manuscript and conducted Western blotting protein assays. EPS helped with the manuscript corrections. AHO helped with HUVEC experiments. ETC and CM performed GSEA assay and analysis. LNSA performed EV mRNA analysis content by qPCR. MSJ helped with IVIS organs-images analysis. SD performed the antibody labeling and helped proofread the manuscript. ECP generated the liposomes and helped to correct the manuscript. YR, AB, and NF helped with reviewing the manuscript and CUBIC and confocal techniques, imaging, and analysis under KMT’s supervision. MP critically reviewed the manuscript and provided the YC-27-Cy5.5 PSMA probe. JG helped develop the idea and extensively and thoroughly reviewed all results, figures, and analysis throughout the experimental as well as the writing phase.

## References

Abid, M.R., S. Guo, T. Minami, K.C. Spokes, K. Ueki, C. Skurk, K. Walsh, and W.C. Aird. 2004. Vascular endothelial growth factor activates PI3K/Akt/forkhead signaling in endothelial cells. Arterioscler Thromb Vasc Biol 24:294–300.

Acedo, S.C., S. Gambero, F.G. Cunha, I. Lorand-Metze, and A. Gambero. 2013. Participation of leptin in the determination of the macrophage phenotype: an additional role in adipocyte and macrophage crosstalk. In Vitro Cell Dev Biol Anim 49:473–478.

Adini, A., T. Kornaga, F. Firoozbakht, and L.E. Benjamin. 2002. Placental growth factor is a survival factor for tumor endothelial cells and macrophages. Cancer Res 62:2749–2752.

Ailuno, G., S. Baldassari, F. Lai, T. Florio, and G. Caviglioli. 2020. Exosomes and Extracellular Vesicles as Emerging Theranostic Platforms in Cancer Research. Cells 9:

Antonarakis, E.S., and M.A. Carducci. 2012. Targeting angiogenesis for the treatment of prostate cancer. Expert Opin Ther Targets 16:365–376.

Barrientos, S., O. Stojadinovic, M.S. Golinko, H. Brem, and M. Tomic-Canic. 2008. Growth factors and cytokines in wound healing. Wound Repair Regen 16:585–601.

Beit-Yannai, E., S. Tabak, and W.D. Stamer. 2018. Physical exosome:exosome interactions. J Cell Mol Med 22:2001–2006.

Biggs, C.N., K.M. Siddiqui, A.A. Al-Zahrani, S. Pardhan, S.I. Brett, Q.Q. Guo, J. Yang, P. Wolf, N.E. Power, P.N. Durfee, C.D. MacMillan, J.L. Townson, J.C. Brinker, N.E. Fleshner, J.I. Izawa, A.F. Chambers, J.L. Chin, and H.S. Leong. 2016. Prostate extracellular vesicles in patient plasma as a liquid biopsy platform for prostate cancer using nanoscale flow cytometry. Oncotarget 7:8839–8849.

Bordas, M., G. Genard, S. Ohl, M. Nessling, K. Richter, T. Roider, S. Dietrich, K.K. Maaß, and M. Seiffert. 2020. Optimized Protocol for Isolation of Small Extracellular Vesicles from Human and Murine Lymphoid Tissues. Int J Mol Sci 21:

Brakenhielm, E., J.B. Burton, M. Johnson, N. Chavarria, K. Morizono, I. Chen, K. Alitalo, and L. Wu. 2007. Modulating metastasis by a lymphangiogenic switch in prostate cancer. Int J Cancer 121:2153–2161.

Bruno, S., C. Grange, M.C. Deregibus, R.A. Calogero, S. Saviozzi, F. Collino, L. Morando, A. Busca, M. Falda, B. Bussolati, C. Tetta, and G. Camussi. 2009. Mesenchymal stem cell-derived microvesicles protect against acute tubular injury. J Am Soc Nephrol 20:1053–1067.

Cao, R., M.A. Björndahl, P. Religa, S. Clasper, S. Garvin, D. Galter, B. Meister, F. Ikomi, K. Tritsaris, S. Dissing, T. Ohhashi, D.G. Jackson, and Y. Cao. 2004. PDGF-BB induces intratumoral lymphangiogenesis and promotes lymphatic metastasis. Cancer Cell 6:333–345.

Cao, R., E. Bråkenhielm, R. Pawliuk, D. Wariaro, M.J. Post, E. Wahlberg, P. Leboulch, and Y. Cao. 2003. Angiogenic synergism, vascular stability and improvement of hind-limb ischemia by a combination of PDGF-BB and FGF-2. Nat Med 9:604–613.

Caromile, L.A., K. Dortche, M.M. Rahman, C.L. Grant, C. Stoddard, F.A. Ferrer, and L.H. Shapiro. 2017. PSMA redirects cell survival signaling from the MAPK to the PI3K-AKT pathways to promote the progression of prostate cancer. Sci Signal 10:

Chen, X., H. Cheng, T. Pan, Y. Liu, Y. Su, C. Ren, D. Huang, X. Zha, and C. Liang. 2015. mTOR regulate EMT through RhoA and Rac1 pathway in prostate cancer. Mol Carcinog 54:1086–1095.

Chen, Y., S. Dhara, S.R. Banerjee, Y. Byun, M. Pullambhatla, R.C. Mease, and M.G. Pomper. 2009. A low molecular weight PSMA-based fluorescent imaging agent for cancer. Biochem Biophys Res Commun 390:624–629.

Corliss, B.A., M.S. Azimi, J.M. Munson, S.M. Peirce, and W.L. Murfee. 2016. Macrophages: An Inflammatory Link Between Angiogenesis and Lymphangiogenesis. Microcirculation 23:95–121.

Costantini, I., J.P. Ghobril, A.P. Di Giovanna, A.L. Allegra Mascaro, L. Silvestri, M.C. Müllenbroich, L. Onofri, V. Conti, F. Vanzi, L. Sacconi, R. Guerrini, H. Markram, G. Iannello, and F.S. Pavone. 2015. A versatile clearing agent for multi-modal brain imaging. Sci Rep 5:9808.

D’Souza-Schorey, C., and J.W. Clancy. 2012. Tumor-derived microvesicles: shedding light on novel microenvironment modulators and prospective cancer biomarkers. Genes Dev 26:1287–1299.

Danaei, M., M. Dehghankhold, S. Ataei, F. Hasanzadeh Davarani, R. Javanmard, A. Dokhani, S. Khorasani, and M.R. Mozafari. 2018. Impact of Particle Size and Polydispersity Index on the Clinical Applications of Lipidic Nanocarrier Systems. Pharmaceutics 10:

Datta, K., M. Muders, H. Zhang, and D.J. Tindall. 2010. Mechanism of lymph node metastasis in prostate cancer. Future Oncol 6:823–836.

De Benedetti, A., and J.R. Graff. 2004. eIF-4E expression and its role in malignancies and metastases. Oncogene 23:3189–3199.

Delprat, V., and C. Michiels. 2021. A bi-directional dialog between vascular cells and monocytes/macrophages regulates tumor progression. Cancer Metastasis Rev 40:477–500.

Di Vizio, D., M. Morello, A.C. Dudley, P.W. Schow, R.M. Adam, S. Morley, D. Mulholland, M. Rotinen, M.H. Hager, L. Insabato, M.A. Moses, F. Demichelis, M.P. Lisanti, H. Wu, M. Klagsbrun, N.A. Bhowmick, M.A. Rubin, C. D’Souza-Schorey, and M.R. Freeman. 2012. Large oncosomes in human prostate cancer tissues and in the circulation of mice with metastatic disease. Am J Pathol 181:1573–1584.

Dikov, D., S. Bachurska, D. Staikov, and V. Sarafian. 2015. Intraepithelial lymphocytes in relation to NIH category IV prostatitis in autopsy prostate. Prostate 75:1074–1084.

Duque, J.L., K.R. Loughlin, R.M. Adam, P.W. Kantoff, D. Zurakowski, and M.R. Freeman. 1999. Plasma levels of vascular endothelial growth factor are increased in patients with metastatic prostate cancer. Urology 54:523–527.

Ferrara, N. 2002. VEGF and the quest for tumour angiogenesis factors. Nat Rev Cancer 2:795–803.

Foster, D.S., R.E. Jones, R.C. Ransom, M.T. Longaker, and J.A. Norton. 2018. The evolving relationship of wound healing and tumor stroma. JCI Insight 3:

Gaceb, A., I. Özen, T. Padel, M. Barbariga, and G. Paul. 2018. Pericytes secrete pro-regenerative molecules in response to platelet-derived growth factor-BB. J Cereb Blood Flow Metab 38:45–57.

García-Silva, S., A. Benito-Martín, L. Nogués, A. Hernández-Barranco, M.S. Mazariegos, V. Santos, M. Hergueta-Redondo, P. Ximénez-Embún, R.P. Kataru, A.A. Lopez, C. Merino, S. Sánchez-Redondo, O. Graña-Castro, I. Matei, J.Á. Nicolás-Avila, R. Torres-Ruiz, S. Rodríguez-Perales, L. Martínez, M. Pérez-Martínez, G. Mata, A. Szumera-Ciećkiewicz, I. Kalinowska, A. Saltari, J.M. Martínez-Gómez, S.A. Hogan, H.U. Saragovi, S. Ortega, C. Garcia-Martin, J. Boskovic, M.P. Levesque, P. Rutkowski, A. Hidalgo, J. Muñoz, D. Megías, B.J. Mehrara, D. Lyden, and H. Peinado. 2021. Melanoma-derived small extracellular vesicles induce lymphangiogenesis and metastasis through an NGFR-dependent mechanism. Nature Cancer

Ghosh, A.K., C.R. Secreto, T.R. Knox, W. Ding, D. Mukhopadhyay, and N.E. Kay. 2010. Circulating microvesicles in B-cell chronic lymphocytic leukemia can stimulate marrow stromal cells: implications for disease progression. Blood 115:1755–1764.

Graff, J.R., B.W. Konicek, R.L. Lynch, C.A. Dumstorf, M.S. Dowless, A.M. McNulty, S.H. Parsons, L.H. Brail, B.M. Colligan, J.W. Koop, B.M. Hurst, J.A. Deddens, B.L. Neubauer, L.F. Stancato, H.W. Carter, L.E. Douglass, and J.H. Carter. 2009. eIF4E activation is commonly elevated in advanced human prostate cancers and significantly related to reduced patient survival. Cancer Res 69:3866–3873.

Grant, C.L., L.A. Caromile, V. Ho, K. Durrani, M.M. Rahman, K.P. Claffey, G.H. Fong, and L.H. Shapiro. 2012. Prostate specific membrane antigen (PSMA) regulates angiogenesis independently of VEGF during ocular neovascularization. PLoS One 7:e41285.

Gupta, D., X. Liang, S. Pavlova, O.P.B. Wiklander, G. Corso, Y. Zhao, O. Saher, J. Bost, A.M. Zickler, A. Piffko, C.L. Maire, F.L. Ricklefs, O. Gustafsson, V.C. Llorente, M.O. Gustafsson, R.B. Bostancioglu, D.R. Mamand, D.W. Hagey, A. Görgens, J.Z. Nordin, and S. El Andaloussi. 2020. Quantification of extracellular vesicles. J Extracell Vesicles 9:1800222.

Haedicke, K., L. Agemy, M. Omar, A. Berezhnoi, S. Roberts, C. Longo-Machado, M. Skubal, K. Nagar, H.T. Hsu, K. Kim, T. Reiner, J. Coleman, V. Ntziachristos, A. Scherz, and J. Grimm. 2020. High-resolution optoacoustic imaging of tissue responses to vascular-targeted therapies. Nat Biomed Eng 4:286–297.

Hirakawa, S., S. Kodama, R. Kunstfeld, K. Kajiya, L.F. Brown, and M. Detmar. 2005. VEGF-A induces tumor and sentinel lymph node lymphangiogenesis and promotes lymphatic metastasis. J Exp Med 201:1089–1099.

Hupe, M.C., C. Philippi, D. Roth, C. Kümpers, J. Ribbat-Idel, F. Becker, V. Joerg, S. Duensing, V.H. Lubczyk, J. Kirfel, V. Sailer, R. Kuefer, A.S. Merseburger, S. Perner, and A. Offermann. 2018. Expression of Prostate-Specific Membrane Antigen (PSMA) on Biopsies Is an Independent Risk Stratifier of Prostate Cancer Patients at Time of Initial Diagnosis. Front Oncol 8:623.

Huth, H.W., D.M. Santos, H.D. Gravina, J.M. Resende, A.M. Goes, M.E. de Lima, and C. Ropert. 2017. Upregulation of p38 pathway accelerates proliferation and migration of MDA-MB-231 breast cancer cells. Oncol Rep 37:2497–2505.

Jansen, B.H.E., Y.J.L. Bodar, G.J.C. Zwezerijnen, D. Meijer, J.P. van der Voorn, J.A. Nieuwenhuijzen, M. Wondergem, T.A. Roeleveld, R. Boellaard, O.S. Hoekstra, R.J.A. van Moorselaar, D.E. Oprea-Lager, and A.N. Vis. 2021. Pelvic lymph-node staging with. Eur J Nucl Med Mol Imaging 48:509–520.

Joncas, F.H., F. Lucien, M. Rouleau, F. Morin, H.S. Leong, F. Pouliot, Y. Fradet, C. Gilbert, and P. Toren. 2019. Plasma extracellular vesicles as phenotypic biomarkers in prostate cancer patients. Prostate 79:1767–1776.

Jorfi, S., E.A. Ansa-Addo, S. Kholia, D. Stratton, S. Valley, S. Lange, and J. Inal. 2015. Inhibition of microvesiculation sensitizes prostate cancer cells to chemotherapy and reduces docetaxel dose required to limit tumor growth in vivo. Sci Rep 5:13006.

Kaittanis, C., C. Andreou, H. Hieronymus, N. Mao, C.A. Foss, M. Eiber, G. Weirich, P. Panchal, A. Gopalan, J. Zurita, S. Achilefu, G. Chiosis, V. Ponomarev, M. Schwaiger, B.S. Carver, M.G. Pomper, and J. Grimm. 2018. Prostate-specific membrane antigen cleavage of vitamin B9 stimulates oncogenic signaling through metabotropic glutamate receptors. J Exp Med 215:159–175.

Kamel, M.G., S. Istanbuly, F.A. Abd-Elhay, M.Y.F. Mohamed, L. Huu-Hoai, M. Sadik, M. Dibas, and N.T. Huy. 2020. Examined and Positive Lymph Node Counts Are Associated with Mortality in Prostate Cancer: A Population-Based Analysis. Urol Int 104:699–709.

Kowal, J., G. Arras, M. Colombo, M. Jouve, J.P. Morath, B. Primdal-Bengtson, F. Dingli, D. Loew, M. Tkach, and C. Théry. 2016. Proteomic comparison defines novel markers to characterize heterogeneous populations of extracellular vesicle subtypes. Proc Natl Acad Sci U S A 113:E968–977.

Krishn, S.R., A. Singh, N. Bowler, A.N. Duffy, A. Friedman, C. Fedele, S. Kurtoglu, S.K. Tripathi, K. Wang, A. Hawkins, A. Sayeed, C.P. Goswami, M.L. Thakur, R.V. Iozzo, S.C. Peiper, W.K. Kelly, and L.R. Languino. 2019. Prostate cancer sheds the αvβ3 integrin in vivo through exosomes. Matrix Biol 77:41–57.

Kuroda, K., A. Horiguchi, T. Asano, and M. Hayakawa. 2008. Prediction of lymphatic invasion by peritumoral lymphatic vessel density in prostate biopsy cores. Prostate 68:1057–1063.

Lai, C.P., O. Mardini, M. Ericsson, S. Prabhakar, C. Maguire, J.W. Chen, B.A. Tannous, and X.O. Breakefield. 2014. Dynamic biodistribution of extracellular vesicles in vivo using a multimodal imaging reporter. ACS Nano 8:483–494.

Lewis, J.S., R.J. Landers, J.C. Underwood, A.L. Harris, and C.E. Lewis. 2000. Expression of vascular endothelial growth factor by macrophages is up-regulated in poorly vascularized areas of breast carcinomas. J Pathol 192:150–158.

Li, W., R.R. Liang, C. Zhou, M.Y. Wu, L. Lian, G.F. Yuan, M.Y. Wang, X. Xie, L.M. Shou, F.R. Gong, K. Chen, W.M. Duan, and M. Tao. 2015. The association between expressions of Ras and CD68 in the angiogenesis of breast cancers. Cancer Cell Int 15:17.

Lilis, I., I. Giopanou, H. Papadaki, and K. Gyftopoulos. 2018. The expression of p-mTOR and COUP-TFII correlates with increased lymphangiogenesis and lymph node metastasis in prostate adenocarcinoma. Urol Oncol 36:311.e327-311.e335.

Linde, N., M. Casanova-Acebes, M.S. Sosa, A. Mortha, A. Rahman, E. Farias, K. Harper, E. Tardio, I. Reyes Torres, J. Jones, J. Condeelis, M. Merad, and J.A. Aguirre-Ghiso. 2018. Macrophages orchestrate breast cancer early dissemination and metastasis. Nat Commun 9:21.

Liu, T., D.E. Mendes, and C.E. Berkman. 2014. Functional prostate-specific membrane antigen is enriched in exosomes from prostate cancer cells. Int J Oncol 44:918–922.

Liu, Z.Q., J.M. Fang, Y.Y. Xiao, Y. Zhao, R. Cui, F. Hu, and Q. Xu. 2015. Prognostic role of vascular endothelial growth factor in prostate cancer: a systematic review and meta-analysis. Int J Clin Exp Med 8:2289–2298.

Livak, K.J., and T.D. Schmittgen. 2001. Analysis of relative gene expression data using real-time quantitative PCR and the 2(-Delta Delta C(T)) Method. Methods 25:402–408.

Lázaro-Ibáñez, E., M. Neuvonen, M. Takatalo, U. Thanigai Arasu, C. Capasso, V. Cerullo, J.S. Rhim, K. Rilla, M. Yliperttula, and P.R. Siljander. 2017. Metastatic state of parent cells influences the uptake and functionality of prostate cancer cell-derived extracellular vesicles. J Extracell Vesicles 6:1354645.

Minami, K., K. Hiwatashi, S. Ueno, M. Sakoda, S. Iino, H. Okumura, M. Hashiguchi, Y. Kawasaki, H. Kurahara, Y. Mataki, K. Maemura, H. Shinchi, and S. Natsugoe. 2018. Prognostic significance of CD68, CD163 and Folate receptor-β positive macrophages in hepatocellular carcinoma. Exp Ther Med 15:4465–4476.

Minciacchi, V.R., M.R. Freeman, and D. Di Vizio. 2015. Extracellular vesicles in cancer: exosomes, microvesicles and the emerging role of large oncosomes. Semin Cell Dev Biol 40:41–51.

Minciacchi, V.R., A. Zijlstra, M.A. Rubin, and D. Di Vizio. 2017. Extracellular vesicles for liquid biopsy in prostate cancer: where are we and where are we headed? Prostate Cancer Prostatic Dis 20:251–258.

Mitchell, P.J., J. Welton, J. Staffurth, J. Court, M.D. Mason, Z. Tabi, and A. Clayton. 2009. Can urinary exosomes act as treatment response markers in prostate cancer? J Transl Med 7:4.

Nojima, S., E.A. Susaki, K. Yoshida, H. Takemoto, N. Tsujimura, S. Iijima, K. Takachi, Y. Nakahara, S. Tahara, K. Ohshima, M. Kurashige, Y. Hori, N. Wada, J.I. Ikeda, A. Kumanogoh, E. Morii, and H.R. Ueda. 2017. CUBIC pathology: three-dimensional imaging for pathological diagnosis. Scientific reports 7:9269.

Nomura, N., S. Pastorino, P. Jiang, G. Lambert, J.R. Crawford, M. Gymnopoulos, D. Piccioni, T. Juarez, S.C. Pingle, M. Makale, and S. Kesari. 2014. Prostate specific membrane antigen (PSMA) expression in primary gliomas and breast cancer brain metastases. Cancer Cell Int 14:26.

Notarangelo, M., C. Zucal, A. Modelska, I. Pesce, G. Scarduelli, C. Potrich, L. Lunelli, C. Pederzolli, P. Pavan, G. la Marca, L. Pasini, P. Ulivi, H. Beltran, F. Demichelis, A. Provenzani, A. Quattrone, and V.G. D’Agostino. 2019. Ultrasensitive detection of cancer biomarkers by nickel-based isolation of polydisperse extracellular vesicles from blood. EBioMedicine 43:114–126.

Omar, M., J. Gateau, and V. Ntziachristos. 2013. Raster-scan optoacoustic mesoscopy in the 25-125 MHz range. Opt Lett 38:2472–2474.

Padda, R.S., F.K. Deng, S.I. Brett, C.N. Biggs, P.N. Durfee, C.J. Brinker, K.C. Williams, and H.S. Leong. 2019. Nanoscale flow cytometry to distinguish subpopulations of prostate extracellular vesicles in patient plasma. Prostate 79:592–603.

Park, Y.H., H.W. Shin, A.R. Jung, O.S. Kwon, Y.J. Choi, J. Park, and J.Y. Lee. 2016. Prostate-specific extracellular vesicles as a novel biomarker in human prostate cancer. Sci Rep 6:30386.

Phuyal, S., N.P. Hessvik, T. Skotland, K. Sandvig, and A. Llorente. 2014. Regulation of exosome release by glycosphingolipids and flotillins. FEBS J 281:2214–2227.

Picus, J., S. Halabi, W.K. Kelly, N.J. Vogelzang, Y.E. Whang, E.B. Kaplan, W.M. Stadler, E.J. Small, and C.a.L.G. B. 2011. A phase 2 study of estramustine, docetaxel, and bevacizumab in men with castrate-resistant prostate cancer: results from Cancer and Leukemia Group B Study 90006. Cancer 117:526–533.

Pore, N., A.K. Gupta, G.J. Cerniglia, Z. Jiang, E.J. Bernhard, S.M. Evans, C.J. Koch, S.M. Hahn, and A. Maity. 2006. Nelfinavir down-regulates hypoxia-inducible factor 1alpha and VEGF expression and increases tumor oxygenation: implications for radiotherapy. Cancer Res 66:9252–9259.

Pore, N., S. Liu, D.A. Haas-Kogan, D.M. O’Rourke, and A. Maity. 2003. PTEN mutation and epidermal growth factor receptor activation regulate vascular endothelial growth factor (VEGF) mRNA expression in human glioblastoma cells by transactivating the proximal VEGF promoter. Cancer Res 63:236–241.

Rolny, C., I. Nilsson, P. Magnusson, A. Armulik, L. Jakobsson, P. Wentzel, P. Lindblom, J. Norlin, C. Betsholtz, R. Heuchel, M. Welsh, and L. Claesson-Welsh. 2006. Platelet-derived growth factor receptor-beta promotes early endothelial cell differentiation. Blood 108:1877–1886.

Sato, S., and A.M. Weaver. 2018. Extracellular vesicles: important collaborators in cancer progression. Essays Biochem 62:149–163.

Sato, Y., S. Nakajima, N. Shiraga, H. Atsumi, S. Yoshida, T. Koller, G. Gerig, and R. Kikinis. 1998. Three-dimensional multi-scale line filter for segmentation and visualization of curvilinear structures in medical images. Med Image Anal 2:143–168.

Schoppmann, S.F., P. Birner, J. Stöckl, R. Kalt, R. Ullrich, C. Caucig, E. Kriehuber, K. Nagy, K. Alitalo, and D. Kerjaschki. 2002. Tumor-associated macrophages express lymphatic endothelial growth factors and are related to peritumoral lymphangiogenesis. Am J Pathol 161:947–956.

Schulke, N., O.A. Varlamova, G.P. Donovan, D. Ma, J.P. Gardner, D.M. Morrissey, R.R. Arrigale, C. Zhan, A.J. Chodera, K.G. Surowitz, P.J. Maddon, W.D. Heston, and W.C. Olson. 2003. The homodimer of prostate-specific membrane antigen is a functional target for cancer therapy. Proc Natl Acad Sci U S A 100:12590–12595.

Senthil, D., G.G. Choudhury, C. McLaurin, and B.S. Kasinath. 2003. Vascular endothelial growth factor induces protein synthesis in renal epithelial cells: a potential role in diabetic nephropathy. Kidney Int 64:468–479.

Shimura, S., G. Yang, S. Ebara, T.M. Wheeler, A. Frolov, and T.C. Thompson. 2000. Reduced infiltration of tumor-associated macrophages in human prostate cancer: association with cancer progression. Cancer Res 60:5857–5861.

Smyth, T., M. Kullberg, N. Malik, P. Smith-Jones, M.W. Graner, and T.J. Anchordoquy. 2015. Biodistribution and delivery efficiency of unmodified tumor-derived exosomes. J Control Release 199:145–155.

Song, W., D. Yan, T. Wei, Q. Liu, X. Zhou, and J. Liu. 2018. Tumor-derived extracellular vesicles in angiogenesis. Biomed Pharmacother 102:1203–1208.

Susaki, E.A., K. Tainaka, D. Perrin, F. Kishino, T. Tawara, T.M. Watanabe, C. Yokoyama, H. Onoe, M. Eguchi, S. Yamaguchi, T. Abe, H. Kiyonari, Y. Shimizu, A. Miyawaki, H. Yokota, and H.R. Ueda. 2014. Whole-brain imaging with single-cell resolution using chemical cocktails and computational analysis. Cell 157:726–739.

Tang, Z., C. Li, B. Kang, G. Gao, and Z. Zhang. 2017. GEPIA: a web server for cancer and normal gene expression profiling and interactive analyses. Nucleic Acids Res 45:W98–W102.

Taraboletti, G., S. D’Ascenzo, I. Giusti, D. Marchetti, P. Borsotti, D. Millimaggi, R. Giavazzi, A. Pavan, and V. Dolo. 2006. Bioavailability of VEGF in tumor-shed vesicles depends on vesicle burst induced by acidic pH. Neoplasia 8:96–103.

Todorova, D., S. Simoncini, R. Lacroix, F. Sabatier, and F. Dignat-George. 2017. Extracellular Vesicles in Angiogenesis. Circ Res 120:1658–1673.

Turay, D., S. Khan, C.J. Diaz Osterman, M.P. Curtis, B. Khaira, J.W. Neidigh, S. Mirshahidi, C.A. Casiano, and N.R. Wall. 2016. Proteomic Profiling of Serum-Derived Exosomes from Ethnically Diverse Prostate Cancer Patients. Cancer Invest 34:1–11.

Ugorski, M., P. Dziegiel, and J. Suchanski. 2016. Podoplanin - a small glycoprotein with many faces. Am J Cancer Res 6:370–386.

Ungersma, S.E., G. Pacheco, C. Ho, S.F. Yee, J. Ross, N. van Bruggen, F.V. Peale, S. Ross, and R.A. Carano. 2010. Vessel imaging with viable tumor analysis for quantification of tumor angiogenesis. Magn Reson Med 63:1637–1647.

van der Kroef, M., T. Carvalheiro, M. Rossato, F. de Wit, M. Cossu, E. Chouri, C.G.K. Wichers, C.P.J. Bekker, L. Beretta, N. Vazirpanah, E. Trombetta, T.R.D.J. Radstake, and C. Angiolilli. 2020. CXCL4 triggers monocytes and macrophages to produce PDGF-BB, culminating in fibroblast activation: Implications for systemic sclerosis. J Autoimmun 111:102444.

Watanabe, R., M. Maekawa, T. Kiyoi, M. Kurata, N. Miura, T. Kikugawa, S. Higashiyama, and T. Saika. 2021. PSMA-positive membranes secreted from prostate cancer cells have potency to transform vascular endothelial cells into an angiogenic state. Prostate 81:1390–1401.

Weidner, N., P.R. Carroll, J. Flax, W. Blumenfeld, and J. Folkman. 1993. Tumor angiogenesis correlates with metastasis in invasive prostate carcinoma. Am J Pathol 143:401–409.

Wernicke, A.G., S. Varma, E.A. Greenwood, P.J. Christos, K.S. Chao, H. Liu, N.H. Bander, and S.J. Shin. 2014. Prostate-specific membrane antigen expression in tumor-associated vasculature of breast cancers. APMIS 122:482–489.

Yang, Z., J. Shi, J. Xie, Y. Wang, J. Sun, T. Liu, Y. Zhao, X. Zhao, X. Wang, Y. Ma, V. Malkoc, C. Chiang, W. Deng, Y. Chen, Y. Fu, K.J. Kwak, Y. Fan, C. Kang, C. Yin, J. Rhee, P. Bertani, J. Otero, W. Lu, K. Yun, A.S. Lee, W. Jiang, L. Teng, B.Y.S. Kim, and L.J. Lee. 2020. Large-scale generation of functional mRNA-encapsulating exosomes via cellular nanoporation. Nat Biomed Eng 4:69–83.

Yoshioka, Y., Y. Konishi, N. Kosaka, T. Katsuda, T. Kato, and T. Ochiya. 2013. Comparative marker analysis of extracellular vesicles in different human cancer types. J Extracell Vesicles 2:

Zaborowski, M.P., L. Balaj, X.O. Breakefield, and C.P. Lai. 2015. Extracellular Vesicles: Composition, Biological Relevance, and Methods of Study. Bioscience 65:783–797.

Zeng, Y., K. Opeskin, L.G. Horvath, R.L. Sutherland, and E.D. Williams. 2005. Lymphatic vessel density and lymph node metastasis in prostate cancer. Prostate 65:222–230.

Zhan, P., Y.N. Ji, and L.K. Yu. 2013. VEGF is associated with the poor survival of patients with prostate cancer: a meta-analysis. Transl Androl Urol 2:99–105.

Zhou, X., Y. Li, X. Jiang, X. Wang, S. Chen, T. Shen, J. You, H. Lu, H. Liao, Z. Li, and Z. Cheng. 2020. Intra-Individual Comparison of 18F-PSMA-1007 and 18F-FDG PET/CT in the Evaluation of Patients With Prostate Cancer. Front Oncol 10:585213.

Zhou, Y., X. Zhu, H. Cui, J. Shi, G. Yuan, S. Shi, and Y. Hu. 2021. The Role of the VEGF Family in Coronary Heart Disease. Front Cardiovasc Med 8:738325.

