## Supplemental data for "PSMA-bearing extracellular vesicles secreted from prostate cancer convert the microenvironment to a tumor-supporting, pro-angiogenic state"

**Supplementary Data**
**Results and figures**

Our method of EV isolation is adapted from a method to isolate larger EV (D'Souza-Schorey and Clancy, 2012; Di Vizio et al., 2012; Gupta et al., 2020; Jorfi et al., 2015; Kowal et al., 2016; Krishn et al., 2019; Smyth et al., 2015) (Supplementary Fig.1A-D). EV from WT or PSMA overexpressing cells measured on average 298.8±22.21 (WT-EV) nm and 267.6±48.74 nm (PSMA-EV, mean±SD), respectively (Fig.1A and B, the yellow square represents 85% of the EV-isolated). EV isolated from LNCaP, PSMA-expressing cells, measured 271.2±75.1 nm. EV isolated from 22RV1 had a significantly smaller size (22RV-EV, Supplementary Fig. 1C) of 162.0±26.66 nm. Liposomes used as an additional PSMA-negative control for *in vitro* experiments, as a cargo-free EV-like vehicle measured 137±5 nm (Supplementary Fig. 1D and E) (Smyth et al., 2015). EV from PC3WT as well as liposomes were used as control (PSMA-negative) vesicles.

Immunoblotting of total EV protein extraction was used to assess protein content of the various EV types. PSMA-EV and PC3PSMA cells contained both as a monomer and homodimer (Schulke et al., 2003) (Fig. 1C, Supplementary 1N). 22RV1-EV contained only small amounts of the monomer, whereas LNCaP-EV have a more significant amount of PSMA monomer relative to the quantity of beta-actin or flotillin-1 (Phuyal et al., 2014) present in larger EV's (Bordas et al., 2020; Smyth et al., 2015; Yoshioka et al., 2013) (Supplementary Fig. 1F). Both WT and PSMA EV contain markers such as flotillin-1(Phuyal et al., 2014; Smyth et al., 2015; Yoshioka et al., 2013), CD9-tetraspanin, β1-integrin and beta-actin (Fig. 1C).(Bordas et al., 2020; Kowal et al., 2016) To adjust for protein content and different protein expression levels between the cell types, all experiments were calibrated to keep a consistent EV amount by Nanosight® evaluation of each isolated batch. The number of EV derived from PSMA expressing PC3 cells (PSMA-EV, LNCaP-EV and 22RV-EV) or EV derived from PC3WT as non PSMA EV controls (WT-EV) is in the range of $10^{10}$ EV to $10^{11}$ (per mL) *in vitro* or *in vivo* experiments, as this number is consistently documented within the literature to be found in PCa human plasma and serum (Di Vizio et al., 2012; Gupta et al., 2020; Minciacchi et al., 2015; Padda et al., 2019; Sato and Weaver, 2018).

Seeking to investigate EV fusion with the membrane of recipient cells, we incubated cells with equal amounts of EV labeled with fluorescent dye (Fig. 1D, Supplementary Fig. 1G-L). PC3WT cells uptake EV from PSMA expressing cells or not up to 24h later (Fig. 1D) without changes in uptake even up 48h later (Supplementary Fig. 1G-H). PSMA detected on recipient cells (Fig. 1C and 1D, Supplementary Fig. 1G-1M) showed a strong PSMA immunofluorescence signal and co-localization with fluorescent PSMA-EV or LNCaP-EV. LNCaP-EV and 22RV-EV has no detectable PSMA in the immunoblot of extracted proteins (Supplementary Fig.1 N), although LNCaP-EV incubated with PC3WT cells for the same period showed strong PSMA staining compared to liposomes PSMA-free-controls (Supplementary Fig. 1O- 1P).

We next investigated EV uptake by stromal cells. Incubation of fluorescent endothelial cells (HUVEC-RFP) with PSMA-EV, LNCaP-EV, and 22RV-EV revealed a differential PSMA expression among samples. The PSMA-mean fluorescence intensity was higher in HUVECs exposed to PSMA-EV (Supplementary Fig. 2A), with preservation of both monomer and homodimer PSMA-forms (Supplementary Fig. 2B). The wide view fluorescent and confocal observations showed homogeneous and equal labeled-EV distribution among samples (Supplementary Fig. 2C).

Liposomes, 22RV1-EV and LNCaP-EV showed significant hydrodynamic size variations among isolated batches, indicating that these vesicles are more polydisperse [(Danaei et al., 2018; Joncas et al., 2019)]. Polydisperse EV tend to aggregate [(Danaei et al., 2018)] and congregate inside vessels more often, representing a bias for vascular localization [(Beit-Yannai et al., 2018)]. The variation among batches of LNCaP-EV, 22RV-EV and liposomes could therefore alter the physical biodistribution *in vivo* [(Joncas et al., 2019),34].

In addition, we tested the EV uptake by an adherent murine macrophage cell line (RAW264.7) exposed to either WT-EV, 22RV-EV, LNCaP-EV and PSMA-EV. Brief exposure (up to 90 minutes) to PSMA-EV or WT-EV resulted in acutely increased EV uptake compared to exposure to LNCaP-EV or 22RV-EV. PSMA presence in the cells was then confirmed by western blotting. No protein degradation pointed to an EV-mediated PSMA transfer to TAM. These results support the feasibility of functional PSMA transfer from PSMA-EV to tumor and TME recipient cells.

To avoid variations in uptake due to different EV sizes or PSMA expression, we designed the *in vivo* experiments to use nude mice with or without PC3 tumors, which were injected daily with either WT- or PSMA-EV, after small tumors detections for up to 25 days. First, we checked if PSMA-EV or WT-EV injected intravenously could disperse equally in healthy animals. No significant differences in biodistribution for WT-EV and PSMA-EV-NIR-labeled at day 25 in mice free of tumors were found (Supplementary Fig.3A-C). Our results from NIR-labeled-EV corroborated studies with radiolabeled EV derived from PC3 cells (Smyth et al., 2015). Moreover, to test any physiological particularities due to EV-accumulation or features we used the$^{18}$F-FDG tracer as a tissue mice pro-inflammatory marker. There were no differences among both groups to liver, spleen, lungs, kidneys, and bones uptakes. PET-images qualitatively showed that the PSMA-EV-group had five times more $^{18}$F-FDG uptake in pelvic and distant lymph nodes than WT-EV injected in mice free of tumors (Supplementary Fig. 3D and E) at 25 days post daily EV injections. Recently, PCa patients with pelvic and distant lymph nodes positive to $^{18}$F-FDG and PSMA-PET had poor prognosis (Jansen et al., 2021; Zhou et al., 2020). No signs of toxicity in mice, such as changes in body weight, ulcerations, bleeding, or labored breathing was observed up to 25 days observations.

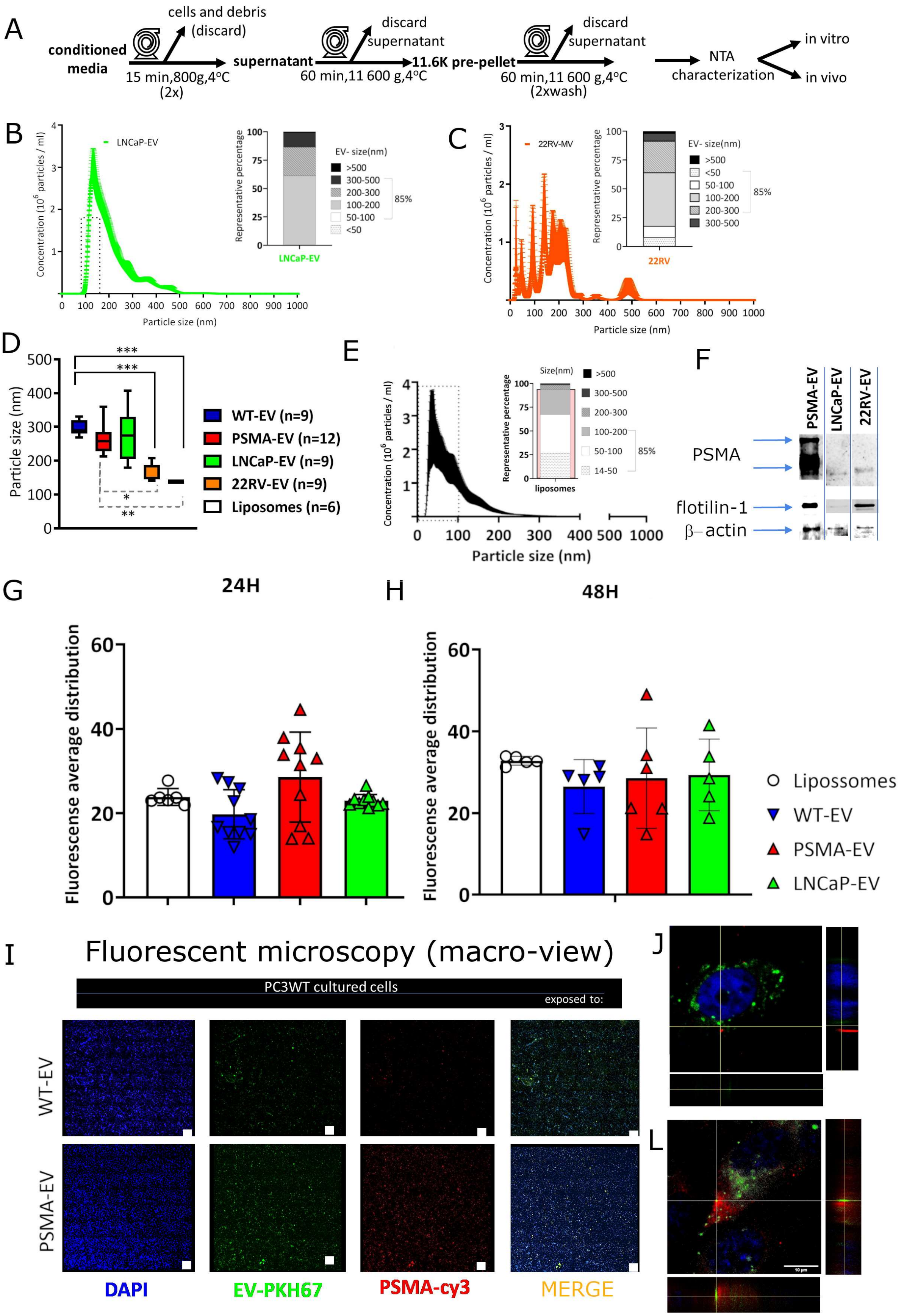
A
conditioned media
cells and debris (discard)
15 min,800g,4°C (2x)
supernatant
discard supernatant
60 min,11 600 g,4°C
11.6K pre-pellet
discard supernatant
60 min,11 600 g,4°C (2xwash)
NTA characterization
in vitro
in vivo
B
LNCaP-EV
Concentration (10^6 particles / ml)
Particle size (nm)
Representative percentage
EV- size(nm)
>500
300-500
200-300
100-200
50-100
<50
85%
C
22RV-MV
22RV
D
Particle size (nm)
WT-EV (n=9)
PSMA-EV (n=12)
LNCaP-EV (n=9)
22RV-EV (n=9)
Liposomes (n=6)
E
Size(nm)
14-50
liposomes
F
PSMA-EV
LNCaP-EV
22RV-EV
PSMA
flotilin-1
β–actin
G
24H
H
48H
Fluorescense average distribution
Lipossomes
WT-EV
PSMA-EV
LNCaP-EV
I
Fluorescent microscopy (macro-view)
PC3WT cultured cells
exposed to:
WT-EV
PSMA-EV
DAPI
EV-PKH67
PSMA-cy3
MERGE
J
L

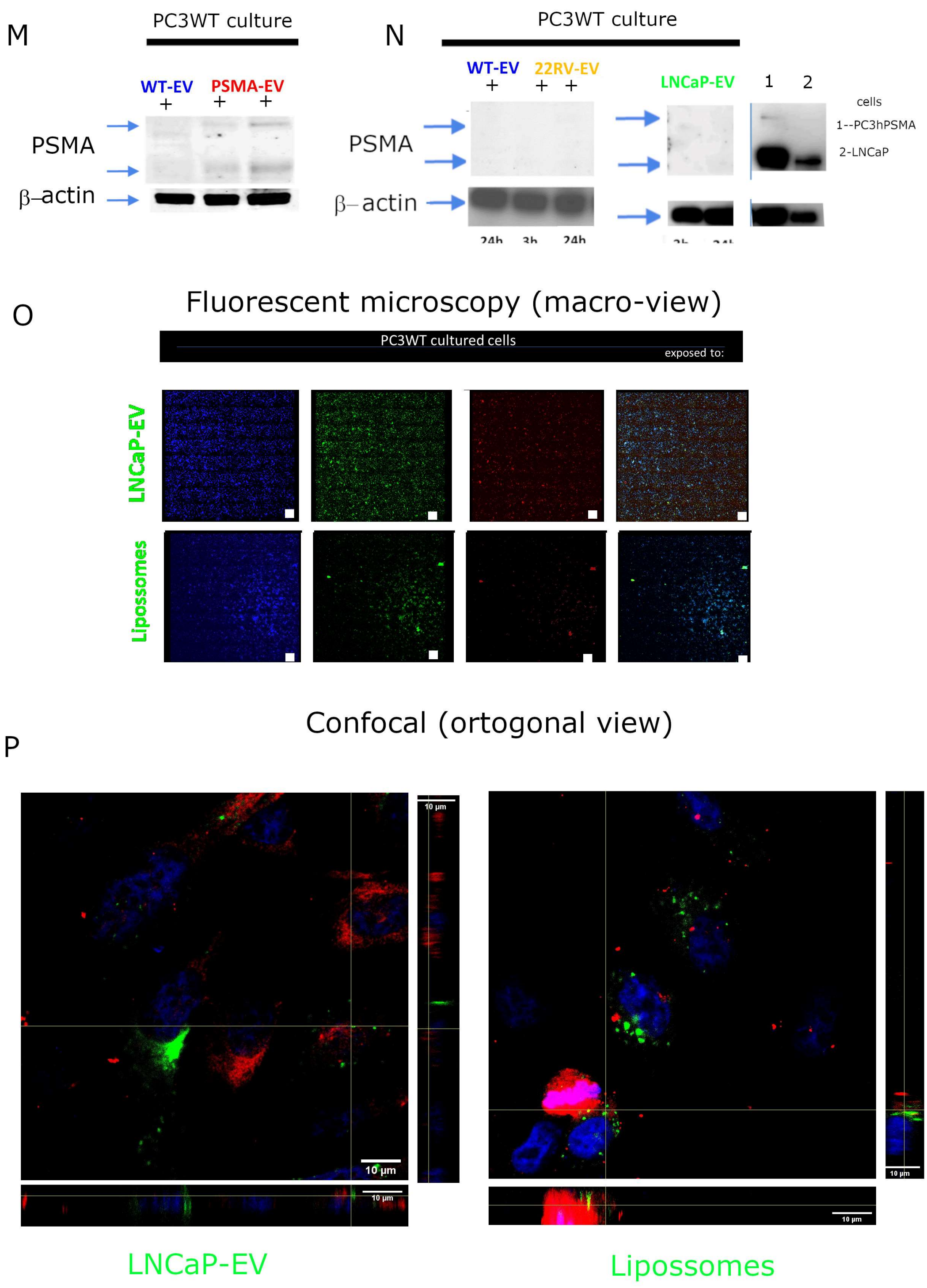


**Supplementary Fig.1:** Tumor-derived extracellular vesicles enriched in microvesicle isolations, physical and biological characterizations. (A) EV isolation scheme based on previous literature. LNCaP and (B) 22RV1 (C) extracellular vesicles. (D) Zetasizer EV size quantitation at 4°C showing from independent isolated batches. P values are: to WT-EV $p<0.0001$, PSMA-EV$p\leq0.002$, 22RV1-EV from WT-EV ($p\leq0.0009$) or PSMA-EV ($p\leq0.0264$). LNCaP-EV and Liposomes (E) characterization by NTA and percentage of each fraction size isolated inside the pellet (insert). (F) PSMA protein expression evaluation by western blot of EV whole protein lysates showing an increased amount of both isoforms of PSMA in PSMA-EV and PSMA-monomer detection in LNCaP-EV or 22RV-EV. Flotillin-1 (a protein marker for larger EV). (G) The mean of fluorescence quantitation of PC3WT cells exposed to fluorescent-PKH67-labeled LNCaP-EV and liposomes up to 24h and 48 (H). The ImageJ is used to quantitate the fluorescence intensity mean of intensity fluorescence in the green channel (These are representative from three independent experiments showing average ± SD). (H) Representative micrographs used for quantitative high-content fluorescence microscopy analysis showing PKH67-labeled-microvesicles uptake after 24h of incubation with recipient-non-expressing PSMA cells (PC3WT) exposed to WT or PSMA-EV. These images are automatically acquired in a wide view matrix (8x8) from a 100x magnification each small square (magnification bar=10um) covering the whole monolayer. (I) PC3WT recipient cells uptake test after 24h to PSMA-EV and WT-EV are taken up by PC3WT cells.(J) Representative confocal micrographs of PC3WT cells after 24 h incubation with PKH67-labeled LNCaP-EV or Liposomes (green) followed by staining with a primary anti-PSMA-antibody and secondary Cy3-antibody (red) and DAPI nuclear staining (blue) (L) Representative micrographs used for quantitative high-content fluorescence microscopy analysis showing PKH67-labeled-LNCaP-EV or Liposomes uptake after 24h of incubation with recipient-non-expressing PSMA cells (PC3WT). These images are automatically acquired in a wide view matrix (8x8) from a 100x magnification each small square (magnification bar=10um). (M-N) The picture shows immunoblotting's of PC3WT cells whole proteins extracts after incubation with (A) 22RV-EV or (B) LNCaP-EV up 24h compared to PSMA expressing cells as a control (PC3hPSMA and LNCaP). (O) Representative PKH67-labeled LNCaP-EV or Liposomes (green) followed by staining with a primary anti-PSMA-antibody and secondary Cy3-antibody (red) and DAPI nuclear staining (blue). These images are automatically acquired in a wide view matrix (8x8) from a 100x magnification each small square (magnification bar=10um) covering the whole monolayer. Confocal micrographs of PC3WT cells after 24 h incubation with without any cell permeabilization (P) EV distributions inside PC3WT cells exposed to LNCaP-EV or Liposomes.

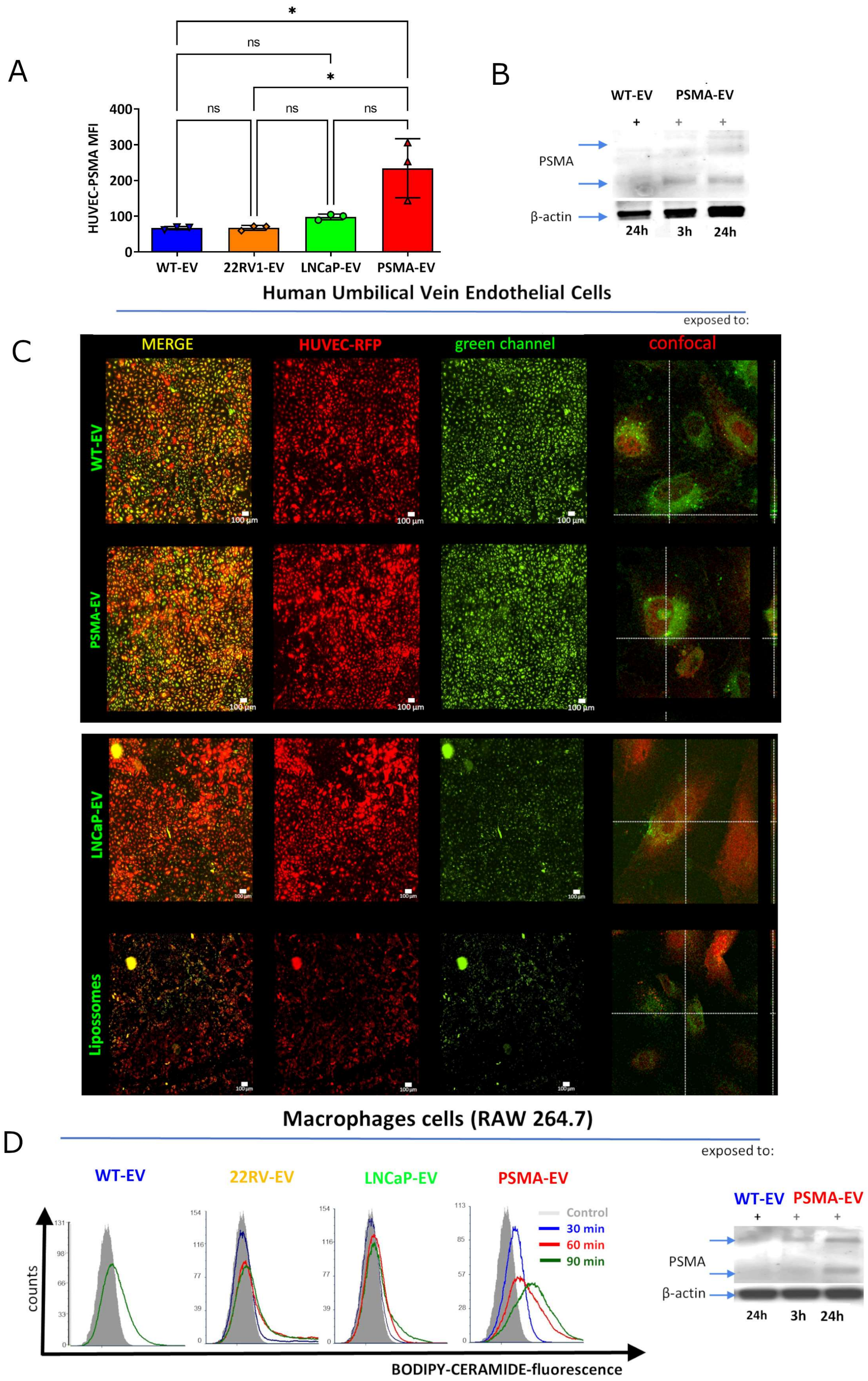


**Supplementary Fig.2:** Vesicles uptake by recipient cells**.** (A) Representative flow cytometry experiments of endothelial cells (HUVEC) incubated with WT-EV, 22RV1-EV, LNCaP-EV, and PSMA-EV and followed by incubation with anti-PSMA antibody conjugated with APC-fluorophore compared to isotype control. (B) Immunoblot of HUVEC whole extracts after incubation with WT-EV and PSMA-EV. (C) Representative confocal micrographs of HUVEC cells after 24 h incubation with PKH67-labeled WT-EV and PSMA-EV, LNCaP-EV or Liposome. Flow cytometry evaluation after labeled-EV incubated with a murine macrophage cell line (RAW264.7, gray filled histogram) for 0.5 (blue), 1 (red), 1.5 (green) hours-time points. PSMA proteins are detected in RAW 264.7 cells whole protein extracts followed by western blotting analysis at 3 or 24h after PSMA-EV exposure.

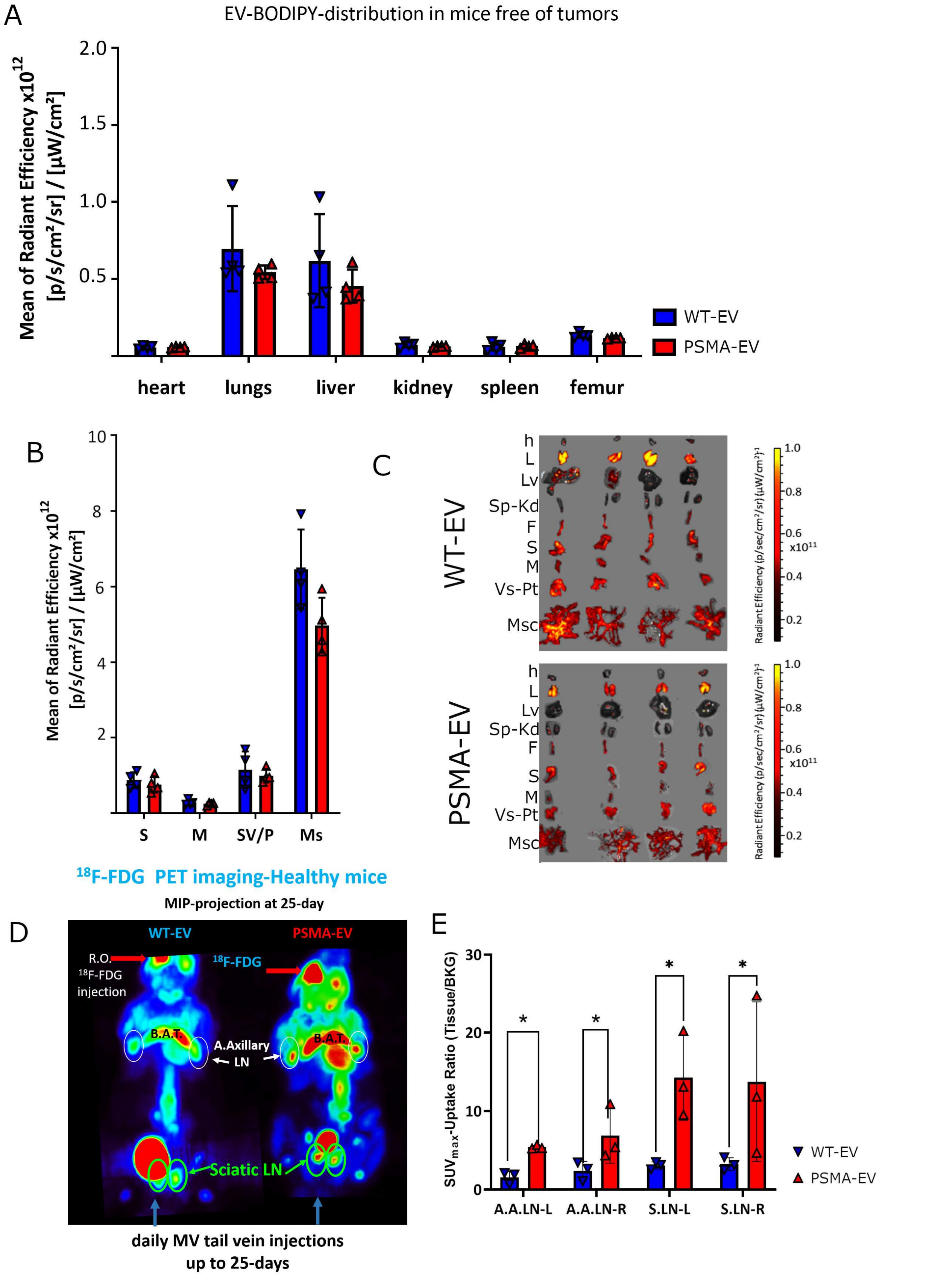


**Supplementary Fig.3:** Representative iVis imaging in mice free of tumors after NIR-EV-labeled (EV-BODIPY-CER) injections up to day 25. (A-B) Quantitation of fluorescent-NIR-EV-labeled-WT or PSMA-EV biodistribution to organs in mice free of tumors. (C) Ex-vivo images of organs imaged by iVis Spectrum .*(h=heart, L=lungs, Lv=Liver, Sp-Kd=Spleen-Kidney, S=stomach, M=muscle, SV-P=Seminal Vesicles-Prostate and MS=Mesocolon).*(D) Same animals observed by PET imaging 1hour after $^{18}$F-FDG biodistribution Each animal received 300uCi tracer injections by retroorbital vein. (E) Note pictoric Lymph nodes increased uptake. Relative quantitation of pelvic and distant lymph nodes. A.A.LN-L and -R: accessory lymph node (LN) left and right; S.LN-L and -R: Sciatic LN left and right.

To determine if PSMA could induce loading of pro-angiogenic factors loading inside extracellular vesicles, we exposed PC3WT cells to exogenous recombinant PSMA (rec-human-PSMA, 0.02µg over 72h), and then isolated EV from these cells. Compared to PSMA-EV and WT-EV, exposure of cells to recombinant PSMA induced an enhancement in pro-angiogenic factors as protein cargo in EV as demonstrated bellow (Supplementary Fig. 4A).

A

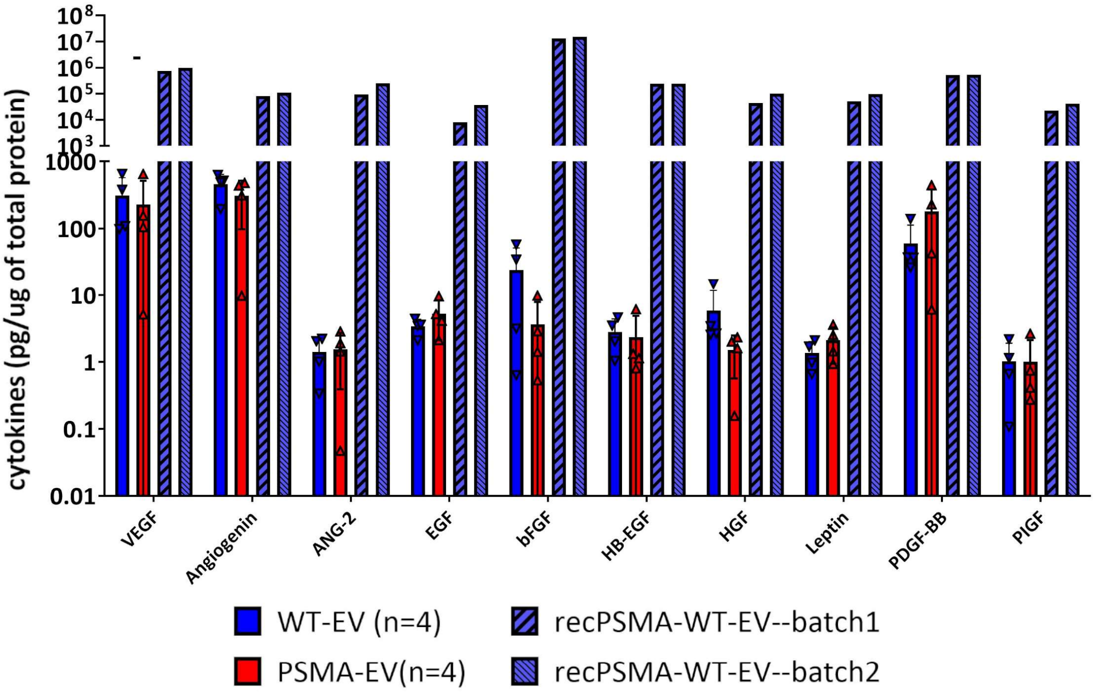


B

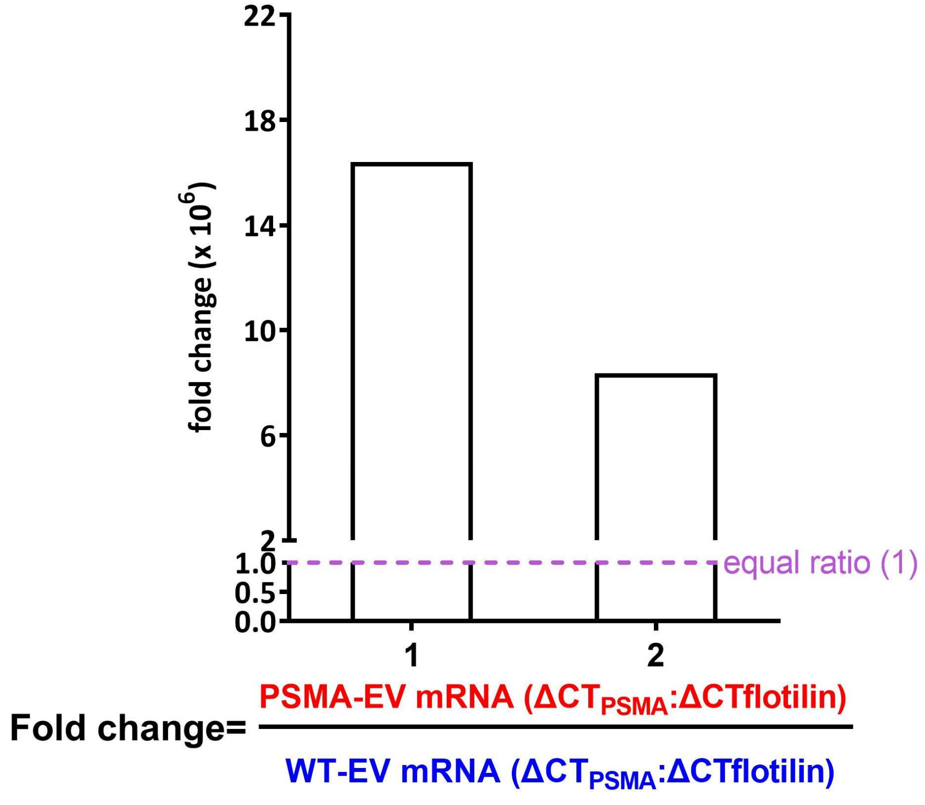


**Supplementary Fig.4: (A)** Extracellular Vesicles protein and mRNA cargo analysis**.** Histogram showing the whole-protein content concentration average (pg/g of protein) and SD of human angiogenesis protein array containing ten different pro-angiogenic factors from 4 independent microvesicles WT-EV or PSMA-EV isolations ($2e^{11}$ microvesicles/each). WT-EV+REC-PSMA were 2-independent vesicles isolations from PC3WT cells exposed to 0.02ug of a recombinant human PSMA/FOLH1 protein (hPSMA) followed by EV extractions and protein cargo analysis ($2e^{11}$ microvesicles/each). WT-EV or PSMA-EV isolations had no differences of protein cargo, although EV isolated from PC3WT cells exposed to hPSMA have enhanced pro-angiogenic protein content inside EV. (angiogenin, vascular endothelial growth factor A- VEGF-A-A, angiopoietin 2-ANG-2, endothelial growth factor-EGF, Basic fibroblast growth factor - bFGF, Heparin-binding EGF-like growth factor-HB-EGF, hepatocyte growth factor -HGF, Leptin, Human Platelet-Derived Growth Factor-BB -PDGF-BB and Placental growth factor-PlGF). **(B)** qRT-PCR for detecting the presence of PSMA mRNA in MV. Flotilin-1 was used as an endogenous control. WT-EV were used as the reference and EV from PSMA overexpressing cells as the target sample. Histogram bars indicate mean. The figure illustrates the fold change between PSMA mRNA levels between EV from 2 independent EV isolations compared to WT-EV to PSMA-EV mRNA ratio levels ($2e^{11}$ microvesicles/each batch and sample).

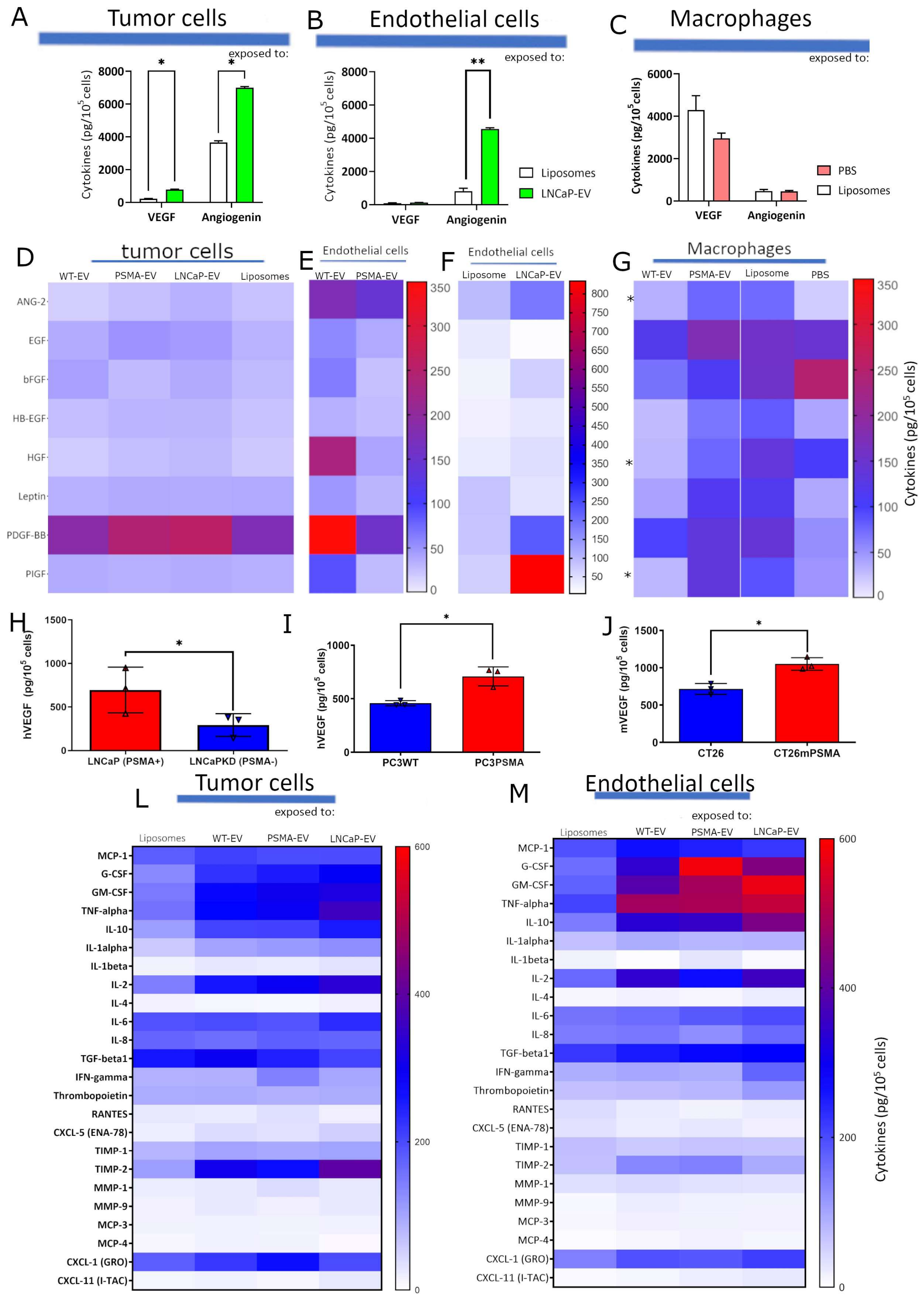


**Supplementary Fig.5:** Histograms and Heatmaps showing the average value of pro-angiogenic factors secretion to cellular media after recipient cells incubations with EV from different cells donors as indicated over the column title. The factors secreted by PC3 cells (2x10$^5$ cell) exposed to EV (2x10$^{10}$) modulates tumor parenchyma (PC3WT) or stroma-like (HUVEC and THP-1) recipient cells *in vitro* (concentration in pg/mL of cells supernatant widening the same number of cells plated). VEGF-A and Angiogenin secretion by (A) tumor, (B) HUVEC and (C) macrophages cells media after cells exposed to Liposomes, LNCaP-EV and PBS. Pro-Angiogenic quantibody array to evaluate factors secretion to media of cells after exposure to EV from different cell donors. (D-G) Significant amounts founded to PlGF secretion by HUVEC (*$p \leq 0.05$) and ANG-2, HGF, and PlGF by Macrophages (*$p \leq 0.03$ and *$p \leq 0.05$).The 2-way ANOVA was used, and p values are represented above. VEGF-A-ELISA detection to ATCC and genetically modified cell lines: (H) LNCaP-PSMA-expressing and LNCaPKD-PSMA-Knockdown; (I) PC3WT and PC3-expressing-PSMA-PC3PSMA; and (J) CT26 and CT26-expressing-murine-PSMA *in vitro*. (D) Murine-PSMA expressing cells (CT26PSMA) model accessed by murine-VEGF-A-ELISA assay normalized by number of cells plated. Data were represented by the average for each group from three independent experiments in triplicates each set with SEM. The t tested (Mann-Whitney) was applied and the p-value was considered significant when **$p \geq 0.01$. (L and M) Heatmap representing the analysis of inflammatory factors that can act as pro-angiogenic stimuli *in vitro*. Data was collected 24 hours after WT-EV or PSMA-EV exposure to tumor parenchyma (PC3WT) or stroma-like (HUVEC) *in vitro*. These data analyzed from the whole-protein media content (average concentration in pg/mL of cells supernatant widening the same number of cells plated) from at least three independent experiments. (Angiopoietin 2 -ANG-2, EGF, Basic fibroblast growth factor - bFGF, Heparin-binding EGF-like growth factor-HB-EGF, hepatocyte growth factor - HGF, Leptin, Human Platelet-Derived Growth Factor-BB -PDGF-BB and Placental growth factor-PLG, PlGF).

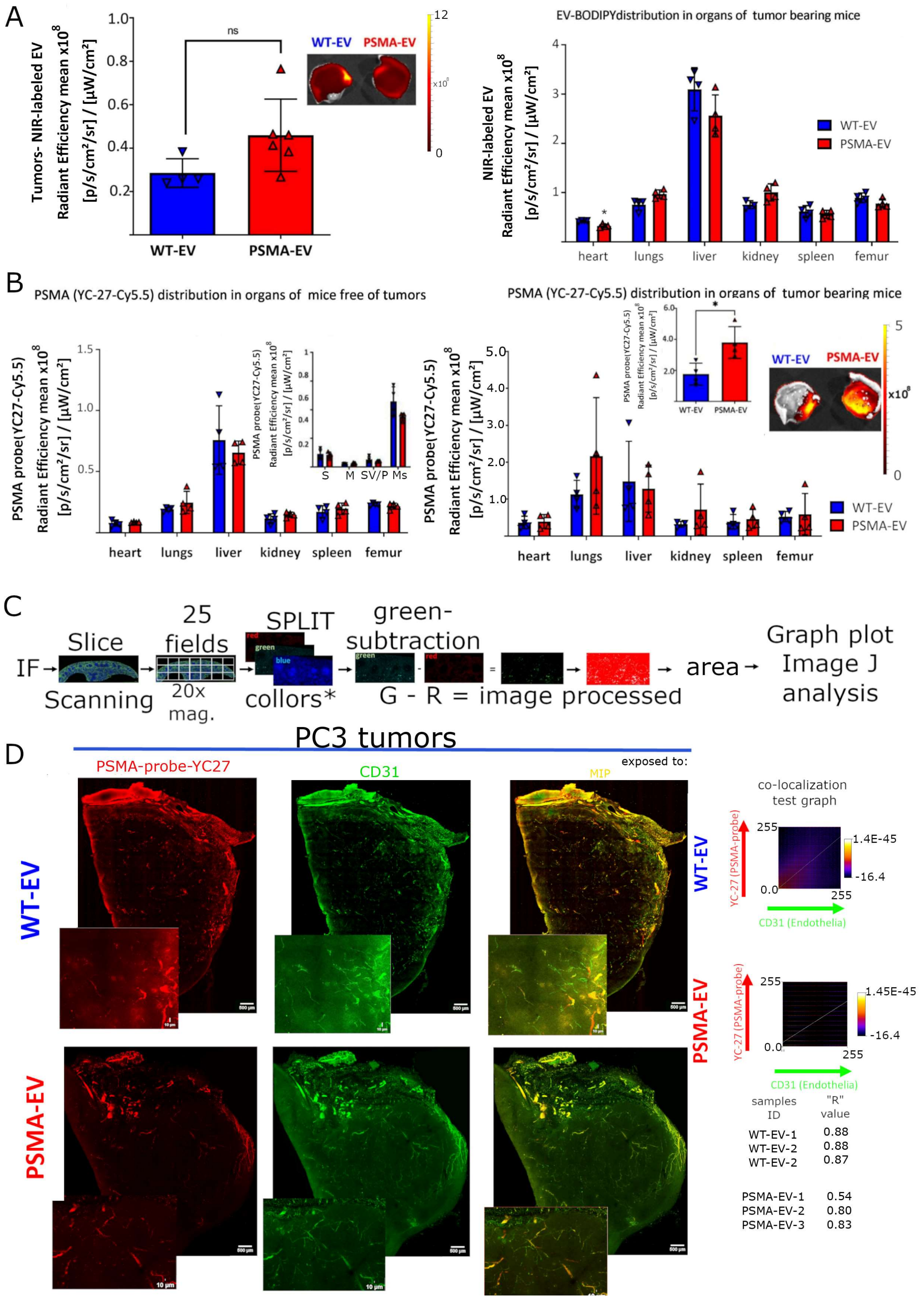


**Supplementary Fig.6:** Representative biodistribution iVis imaging of organs in mice free of tumors or bearing tumors after EV-labeled with near infra-red - BODIPY-CER up to 25-days and *PSMA*

*(YC27-Cy5.5) injected probe* (injected at day 24 and imaged at day 25). Ex-vivo quantitation of NIR-labeled-WT or PSMA-EV biodistribution to tumors (A) organs in mice bearing tumors at day 25 after WT- or PSMA-EV daily tail vein injections. Ex-vivo images and quantitation of *PSMA (YC27-Cy5.5)* distribution (image of 24h) to organs by iVis Spectrum in animals free of tumors (B-left). PSMA-YC37-Cy5.5 distribution (image of 24h) in tumors (insert) and organs from tumor bearing mice exposed to WT- or PSMA-EV at day 25 after daily-EV-injections. (C) Explicative scheme to show the method chosen to quantify immunofluorescences without blood fluorescence interference. (D) PSMA, CD31, and CD68 tumors fluorescence quantitation**.** Confocal Mean-intensity-projection (MIP) of images from half of tumors with 300um thickness. Tissue cleared by CUBIC-solution showing angiogenesis morphology. PC3 xenografts tumors were observed at 25$^{th}$ day after WT-EV (upper) or PSMA-EV (bottom) treatment-daily. CD31+ endothelial cells (green, as described in Materials and Methods) observed by immunofluorescence in combination with a fluorescent agent injected through tail vein at day 24 to show PSMA (YC27-Cy5.5, in red). These experimental settings also allowed the colocalization tests and quantitation of areas with positive CD31 and PSMA as showed at right-column by comparing each pixel spatial-position of the whole MIP image (bottom-left).

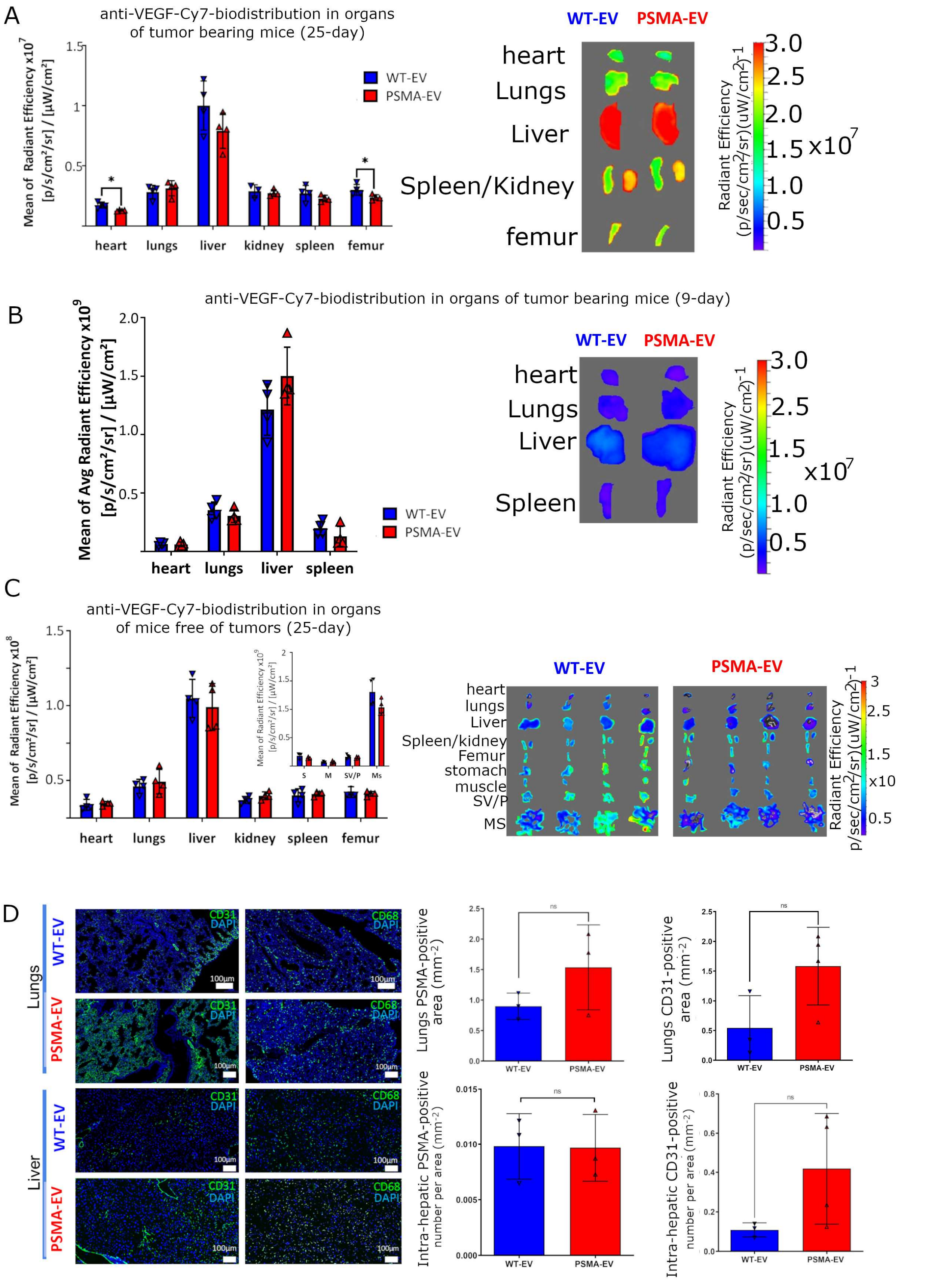


**Supplementary Fig.7:** *Representative anti-VEGF-A-Cy7 biodistribution iVis imaging and quantitation of fluorescence in organs from mice bearing tumors and after EV injections at day 25* ***(A)****; day-9* ***(B)*** *or in organ mice free of tumors* ***(C)****. The images are the merge result between the picture*

*of grayscale organs or tumors and the fluorescent signal is red hot (Bodipy-CER and YC27-Cy5.5) and in Rainbow (VEGF-A) color scale.* ***(D)*** *Lungs and Liver representative images and quantitation showing PSMA, CD31 positive areas with graphs representing one experiment out of 2. (Scale bar=100um). The graphics data represent the average and SD of 4 animals-each. All statistical analysis was conducted by using the unpaired t-test where *p<0.02.(h=heart, L=lungs, Lv=Liver, Spl/Kd=Spleen/Kidney, S=stomach, M=muscle, SV/P=Seminal Vesicles/Prostate and MS=Mesocolon).*

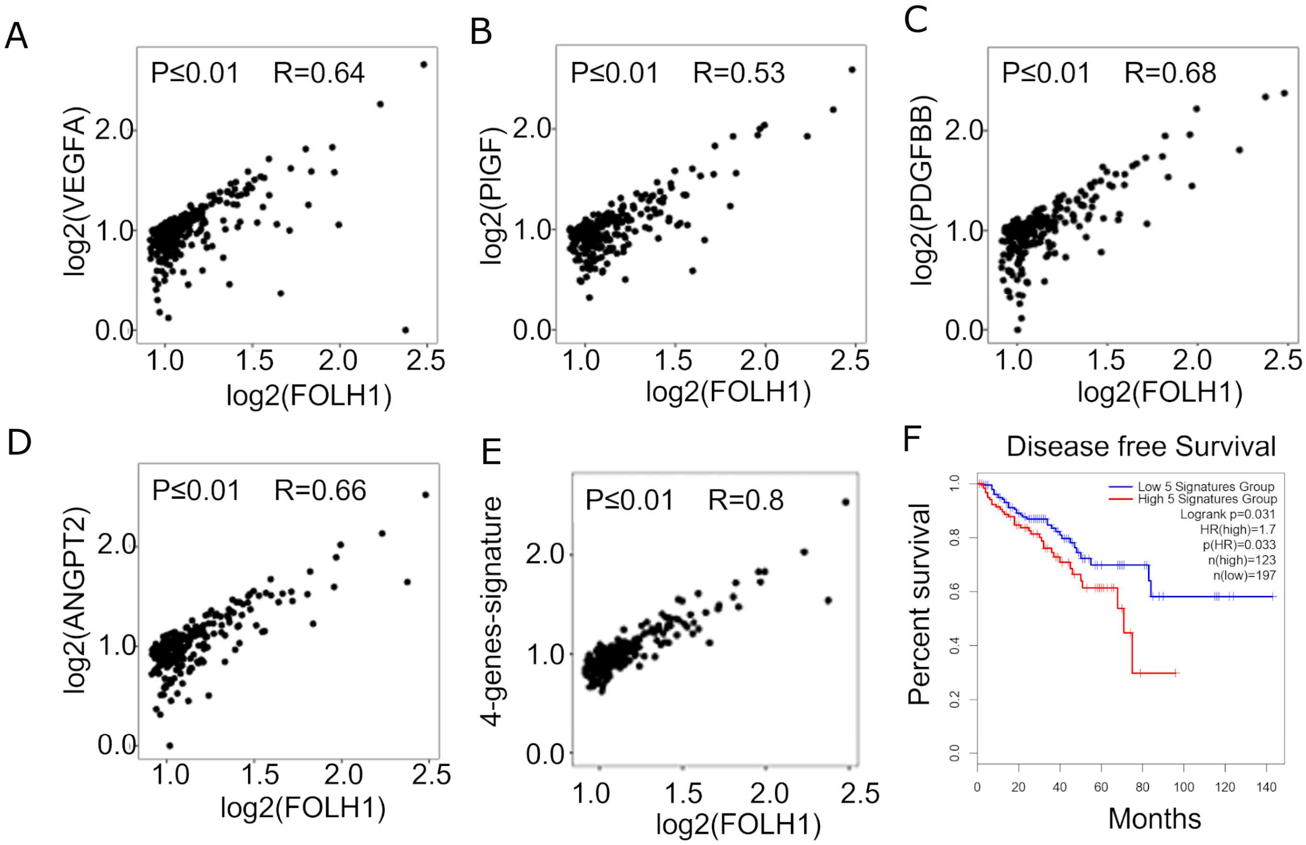


**Supplementary Fig.8**. ***(A-F)*** *Co-expression analysis showing PCa samples with PSMA higher expression and positively correlated to other genes as* ***(A)*** *vascular endothelial growth factor (VEGF-*

*A),* ***(B)*** *placental growth factor (PIGF),* ***(C)*** *human platelet-derived growth factor-B (PDGFB and* ***(D)*** *angiopoietin-2 (ANGPT2) pro-angiogenic genes separately* ***(E)*** *or all together* ***(F)*** *The same patient cohort showing higher PSMA (FOLH1) and proangiogenic gene expression simultaneously analysis versus disease free survival. (GEO # 104419; num(T)=492; linear correlations with "P" and "R" indicated in each section).*

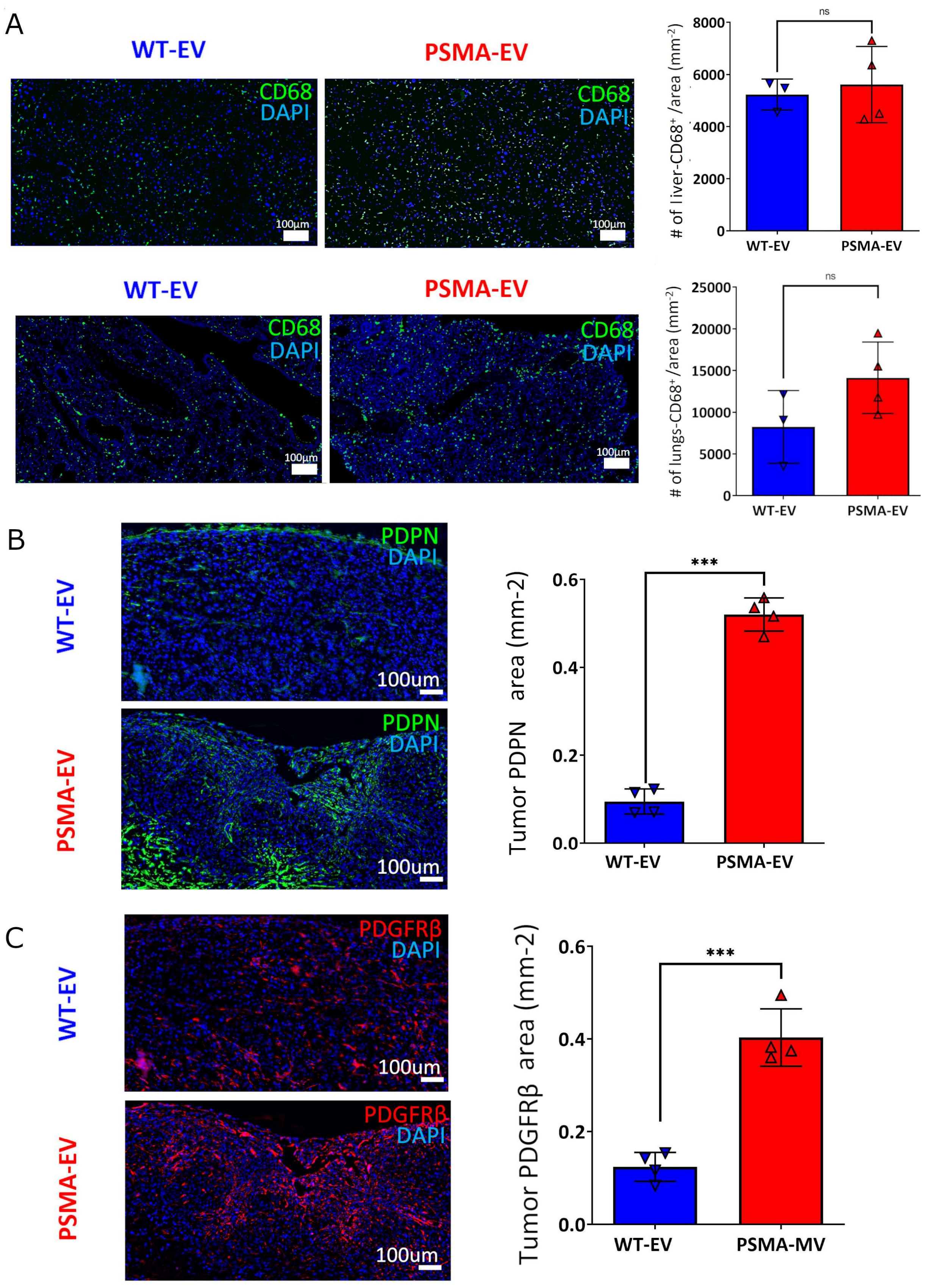


**Supplementary Fig.9:** CD68$^+$, PDPN $^{+,}$ PDGFRβ$^+$ (green, green and red respectively) positive cells quantitation inside tissues collected from mice bearing tumors at day 25, exposed to daily injections with EV. (A) Liver and Lungs representative number of $CD68^+$ cells images and quantitation showing positive areas with graphs representing one experiment out of 2. (Scale bar=100um). The data represents the average and SD of animals each (n=4). All statistical analysis was conducted by using Student's unpaired t-test, and the results were considered significant where *P ≤0.05, ***P<0.0005).

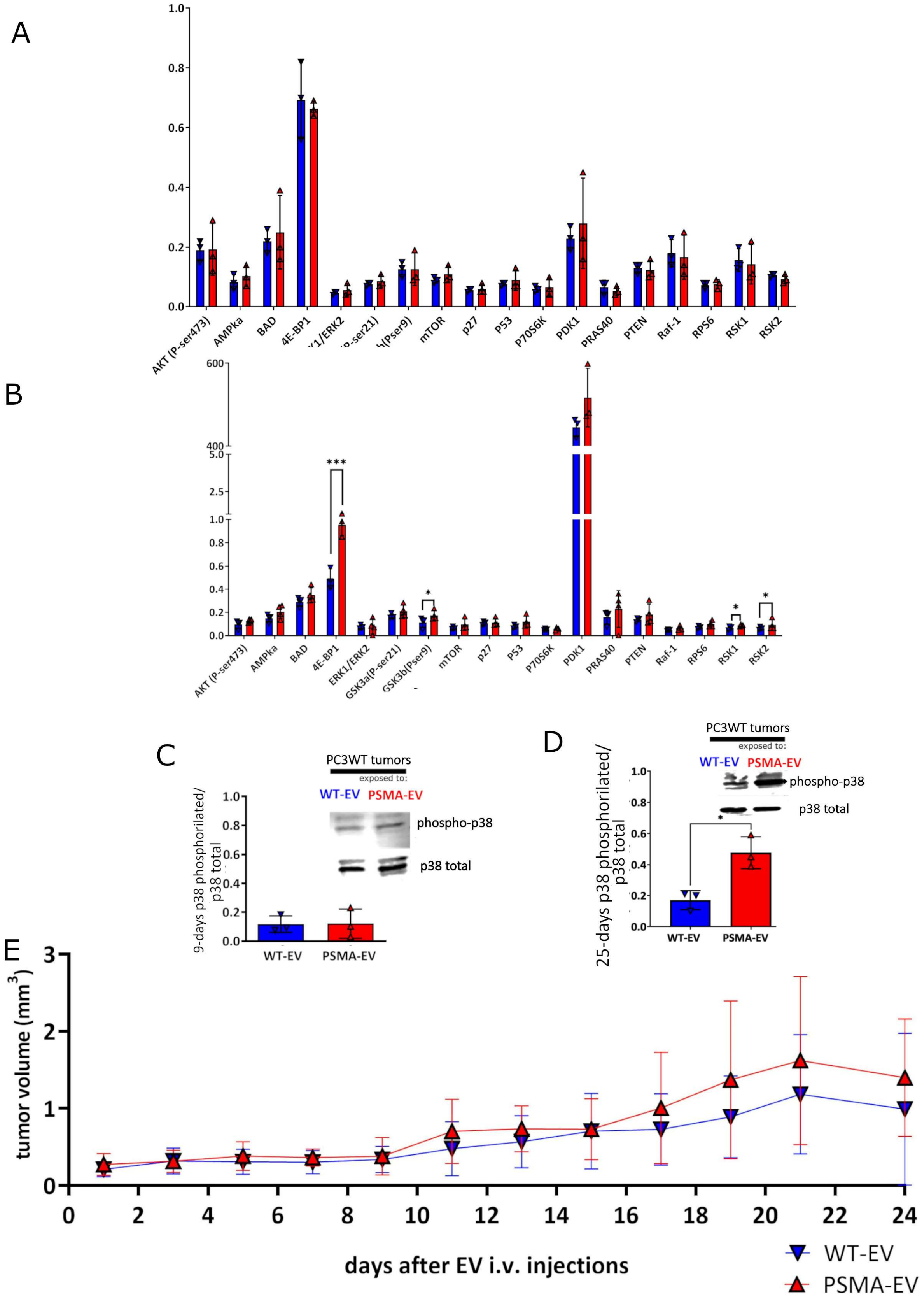


**Supplementary Fig.10 -** *Representative graphics showing:* ***(A and B)*** *Upregulated phosphorylation of PDK-1 and 4EBP-1was observed after 9 or 25 days of treatment with EV. These*

*results were obtained by a multi-protein-AKT-array (AAH-AKT-1-2, RayBiotech) analysis of whole protein extracts from tumors. The factors analyzed were: Akt (P-Ser473), AMPKa (P-THR172), BAD (P-Ser112), 4EBP-1(P-Thr36), ERK1 (P-T202/Y204) / ERK2(P-Y185/Y187), GSK3a (P-Ser21), GSK3b (P-Ser9), mTOR (P-Ser2448), p27 (P-Thr198), P53 (P-Ser15), P70S6K (P-Thr421/Ser424), PDK1 (P-Ser241), PRAS40 (P-Thr246), PTEN (P-Ser380), Raf-1 (Ser301), RPS6 (P-Ser235/236),RSK1 (P-Ser380) and RSK2 (P-Ser386). The statistical analysis was conducted by using the unpaired t-test comparing WT-EV to PSMA-EV means (* p =0.01, ***p=0.0001). The quantitation is representative of 2 independent experiments with the same results. Total extracted protein levels from PC3WT tumors exposed to WT-EV or PSMA-EV on day 9***(C)*** *and 25* ***(D)*** *of EV treatment was analyzed by western blot and semi-quantitative analysis. Total p38 protein was used as a loading control for phosphorylated-p38.* ***(E)*** *Representative tumor volume growth curve.*
